# Pulsations and flows in tissues: two collective dynamics with simple cellular rules

**DOI:** 10.1101/2020.07.29.226357

**Authors:** Raghavan Thiagarajan, Alka Bhat, Guillaume Salbreux, Mandar M. Inamdar, Daniel Riveline

## Abstract

Collective motions of epithelial cells *in vivo* are essential for morphogenesis in developmental biology. Tissues elongate, contract, flow, and oscillate, thus sculpting embryos. These tissue level dynamics are known, but the physical mechanisms at the cellular level are unclear, with various behaviors depending on the tissues and species. Moreover, investigations on *in vitro* tissue behavior usually focus on only one type of cell dynamics and use diverse theoretical approaches, making systematic comparisons between studies challenging. Here, we show that a single epithelial monolayer of Madin Darby Canine Kidney (MDCK) cells can exhibit two types of local tissue kinematics, pulsations and long range coherent flows. We analyzed these distinct motions by using quantitative live imaging. We also report that these motions can be controlled with internal and external cues such as specific inhibitors, and friction modulation of the substrate by microcontact printing method. We further demonstrate with a unified vertex model that both behaviors depend on the competition between velocity alignment and random diffusion of cell polarization. When alignment and diffusion are comparable, a pulsatile flow emerges, whereas the tissue undergoes long-range flows when velocity alignment dominates. We propose that environmental friction, acto-myosin distributions, and cell polarization kinetics are important in regulating the dynamics of tissue morphogenesis.

## Introduction

During development, tissue dynamics shapes organs with remarkable choreography in space and time [1–5]. Groups of hundreds/thousands of cells flow like liquids in developing embryos in *Drosophila* [6], zebrafish [7], and insects [8]. Solid-like pulsatile movements are also reported in *Caenorhabditis elegans* [9]; in the developing heart of zebrafish, where collective oscillatory motion is required for its proper development[10]; and, more generally, in muscle layers spanning epithelial layers [11]. These millimeter-scale changes in shapes involve the acto-myosin cytoskeleton, which drives tissue movement and generates stress at the organism level [1, 3, 12]. These motions and deformations drive the shaping of tissues, eventually establishing organ functions through cell differentiation. Altogether, flows and pulsations are general phenomena in morphogenesis.

Strikingly, these changes in shapes are reiterated in evolution with certain ‘families’ of tissue transformation, such as elongation and gastrulation, among many morphogenetic events. Even if their genetic backgrounds are different, the basic cellular mechanisms are expected to be shared among different biological model systems [13]. Indeed, simple physical rules to achieve morphogenesis are known: (i) tissues are composed of cells that change their shapes over time, (ii) cells in tissues interact with their neighbors by adhering to each other via adherens junctions and to the surrounding extra-cellular matrix through focal contacts [14]. It is puzzling, therefore, that the same monolayer of cells connected to each other and the matrix can undergo distinct dynamics. Regardless, the subtle complexity of cellular interactions that remain to be elucidated result in self-organization at the tissue level and minor perturbations in the program can lead to very different tissue morphogenesis reported *in vivo* [15]. Indeed for decades the existence of cellular flows and pulsations with groups of associated cells has been recognized in developmental biology across model systems.

Various biological systems share a number of dynamical morphogenetic events *in vivo*, but they are rarely investigated with a unified physical model to observe their diversity. For instance, during blood vessel formation, seven events were reported to contribute to angiogenesis, i.e., cell migration, anastomosis, apical membrane fusion, cell elongation and rearrangements, cell splitting, cell self-fusion, and cell division [16, 17]. Each of them may contribute to the formation of the vessel and the associated cellular flows. More generally, this so-called mechanism of ‘branch formation’ is essential to shape organs, and goes beyond the formation of the vascular systems [18, 19]. In fact, during branch formation, the lungs, kidneys, and mammary glands undergo these different dynamic events at the single-cell level, leading to global cellular reorganizations. Cells at the ‘tip’ of the branches are mesenchymal (behaving as migrating single cells) and they are connected to the ‘stalk’ made of epithelial cells (behaving as a connected tissue), which may undergo passive motion and/or rearrangements through neighbor exchanges and cell proliferation for tracheal branching and vertebrate angiogenesis [18, 19]. However, the identification of conserved principles still lacks a proper understanding of physical rules for cells interactions that lead to global movements.

Several types of single-cell dynamics in morphogenesis, such as the growth of protrusions through filopodia and lamellipodia, binding to the surrounding extracellular matrix with focal contacts, and connecting with the neighboring cells through adherens junctions, have been reported [20]. Cells can also have motility either as independent individuals or collectively in a group. In the latter case, it is often difficult to disentangle the motion of cells as arising from, for example, either active single-cell motion or from cell proliferation due to an increase in cell density and the associated reorganization of tissues with gradients in cell density stress [21]. Thus, in the search for quantitative methods to characterize collective motions beyond experiments with live microscopy, new approaches from theoretical physics for active matter are needed [22]. Indeed, tight and quantitative comparisons with numerical simulations are known to be useful in testing these collective dynamics of groups of cells emerging out of rules of interactions at the cellular levels.

Another level of complexity is related to the cell state, which may change during development, for example, through differentiation and mesenchymal-to-epithelial transitions [23]. Changes in cell states can be either chemical or mechanical or both during the subsequent steps following exit from pluripotency. Biological changes in cells are also accompanied by mechanical modifications and alterations in contractility and intercellular adhesion due to changes in protein expression [23, 24]. However, if the chemical state alters the mechanical state of the cells at any time, physical principles governing these cellular dynamics are robust and still at play irrespective of the cell states. Thus, the search for physical principles is essential to decipher the rules of tissue morphogenesis during development. Hence, unifying experiments tightly coupled to models are expected to unravel new means by which cells organize themselves in tissue.

Toward this aim, we use the dynamics of epithelial Madine Darby Canine Kidney (MDCK) monolayer *in vitro* on flat 2D surfaces as a paradigm for morphogenesis in general [25–28]. Collective effects in epithelial monolayers *in vitro* have been reported with new insights into the physics of active matter and developmental biology [29]. The concepts of topological defects and their importance in setting new rules for morphogenesis [30] and the influence of cellular flow patterns on stress distribution and cell fates [31] are just two of the many examples which show the impact of these new ideas. In addition, these concepts have also served as paradigms for new phenomena *in vivo*, for example, in identifying the key function of topological defects in hydra development [32], or determining the principles behind the role of cellular flows and waves *in vivo* in major morphogenetic events in *Drosophila* [33].

In this study, we focused on two dynamics of collective cell movements, pulsations and flows, which co-exist in standard 2D cell culture *in vitro*. We reasoned that the experimental control of each type of motion would guide us in understanding their cellular origins and successfully tested this assumption. First, we generated pulsations by modifying the friction between the tissue with surfaces patterned with fibronectin at the spontaneous coherence length scale of the monolayer. We then induced cellular flows by exposing the monolayer to cytoskeletal drugs. We also generated pulsations in conditions of flows by using laser ablation. To shed light on the underlying cellular mechanisms, we performed numerical simulations based on a unified vertex model with cell motility and successfully reproduced both types of motion by modulating just two factors.

### Characterizing the spontaneous flow in epithelial monolayers

We seeded MDCK cells on glass substrate and acquired time-lapse images after the monolayer reached confluency. We start with a initial seeding concentration of 10^6^ cells over 175 mm^2^ and allow the monolayers to become confluent. As noted previously [26, 27], confluent monolayers of MDCK underwent complex flow patterns that changed rapidly over time and eventually slowed down because of cell proliferation. To quantify the spatio-temporal patterns of cellular movements, we acquired the time-dependent velocity field using particle image velocimetry (PIV) on the time-lapse images (Figures 1a and 1d and Figure S3a). The MDCK monolayer seemed to exhibit temporally periodic patterns of spatially contractile and expanding cellular movements. To quantify local area changes in the tissue due to these pulsatory patterns, we used the velocity field to calculate the flow divergence (see Velocity Field and Correlation Functions section in STAR Methods for details). The divergence and velocity fields showed that the flow field was correlated in space on a characteristic length scale of ≈ 200 *µ*m (Figures 1c and 1f). Focusing on the domains of this size (Figures 1a and 1d), we noticed regions with visible pulses of contracting and expanding tissues (Figures 1a, 1c, S3b and Movie 1) for approximately 5 h, or areas where the flow appeared to be more uniform and correlated over a larger length scale (Figures 1d-1f, S3c and Movie 2). Although the pulsations were changing locations randomly, there were many stable pulsations that did not change their location over the observed dominant period of ≈ 5 h, i.e., the divergence maxima was observed in the same location.

**Figure 1:**
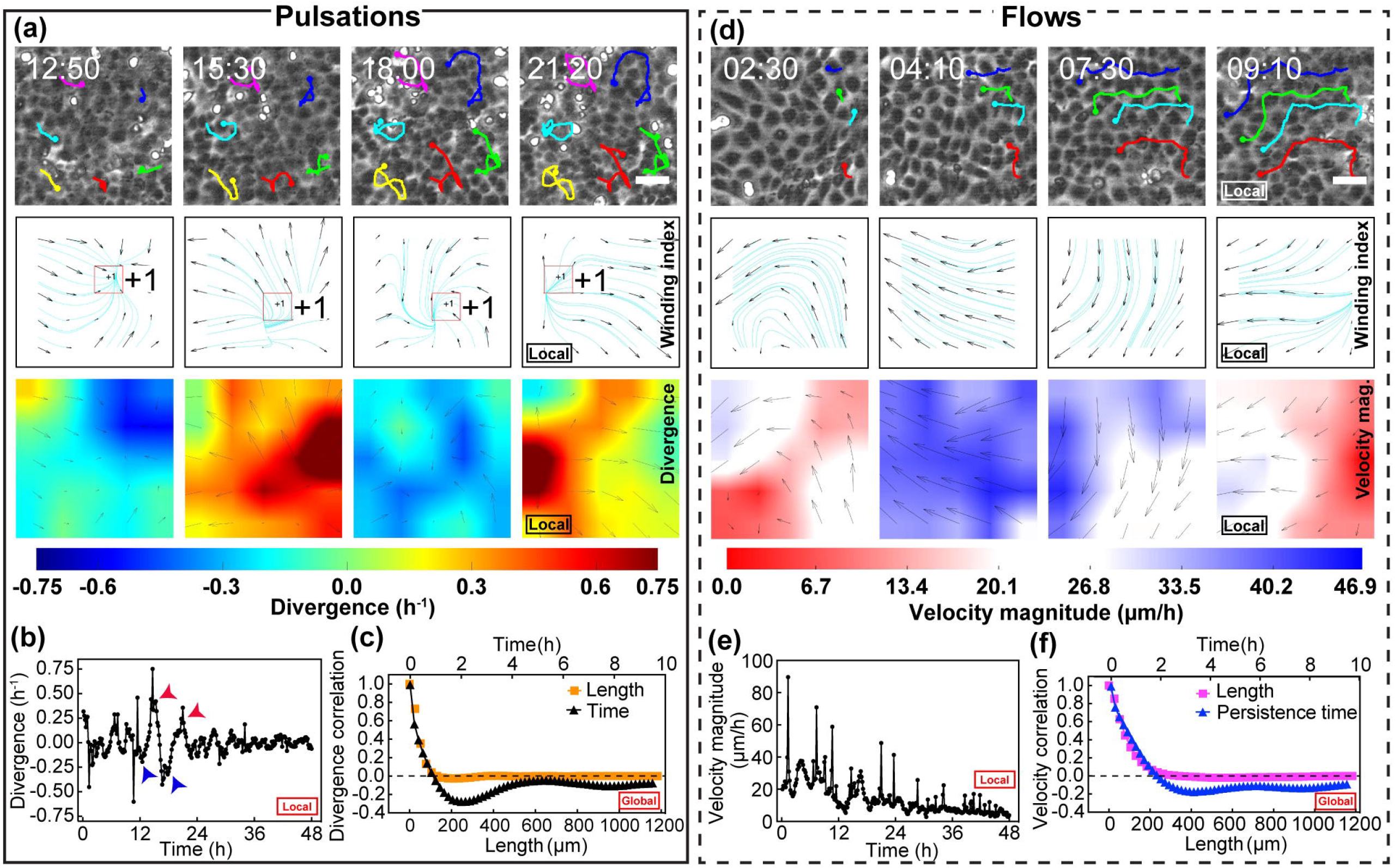
Spontaneous pulsations and flows in an epithelial monolayer. (a), (b), (d), and (e) represent the local characteristics of the monolayer. (a) shows snapshots of the contraction and expansion phases of one period of a pulsating domain, and (d) shows snapshots of flows occurring in the same domain at a different time point. In (a) and (d): the first row corresponds to phase-contrast images, and tracking highlights the back-and-forth movement of cells during pulsation in (a) and flow trajectory in (d); the second row shows the winding number analysis for the snapshots in the first row, and the third row shows the corresponding divergence and velocity magnitude plots, respectively, for (a) and (d). Scale bar 50 *µ*m. Time is in hh:mm. In those panels indicating the nature of the divergence field, blue indicates contraction, and red indicates expansion; similarly, in those panels indicating velocity magnitude, red and blue indicate low and high velocities, respectively. The numerical values are specified in the corresponding color bars. (b) The mean divergence plot for the pulsatile domain over 48 h represented in (a) and in Movie 1. (e) The mean velocity magnitude plot of the flow domain shown in (d) and in Movie 2. Arrows in (b) indicate the data points corresponding to snapshots in (a). (c) and (f) represent the global characteristics of the monolayer and show the divergence and velocity correlations, respectively, for distance and time for the same experiment over 48 h. See also Figures S1 and S3, Movies 1 and 2.

We use the structure of velocity field vis-a-vis streamlines and the presence of ±1 topological defects (see Figures 1a and 1d, second panel and Winding number section in the STAR Methods) in order to distinguish between the two types of tissue kinematics, pulsations and flow, presented in Figures 1a and 1d (first panel), respectively. In the monolayer region with pulsations (Figure 1a), we typically saw 2−4 pronounced pulses in the velocity divergence (see Figure 1b and the autocorrelation plot in Figure S3d). Moreover, the structure of the velocity field is complex, and consequently we observe a number of +1 and −1 defects (Figure 1a, second row). On the other hand, in the monolayer domain with flow (Figure 1d, second panel), the velocity streamlines have a simpler structure resulting in far fewer topological defects. However, we clarify that even in this case, there is an underlying pulsatory velocity pattern (Figure S3c), but it is overwhelmed by the overall flow of the cells. Nonetheless, the coherent flow within the given domain does not imply large-scale directionality since the length scale associated with the velocity is around 200 *µ*m (Figures 1c and 1f), which is chosen to be the approximate size of the domain in Figures 1a and 1d (first panel), and is much smaller than the total field of view of the monolayer (Figure S3a).

The classification of monolayer velocity patterns within the given domains as pulsations or flows can provide a basis for further perturbation experiments. However, to quantify the tissue kinematic behavior over wider length scales, we calculated correlation functions in space and time for the velocity field and its divergence for the entire region under experimental consideration (Figures 1f and 1c, respectively, and Figure S3a). We determined the correlation function of the divergence at the same spatial but different time points, ⟨div(**x**, *t*)div(**x**, *t* + *τ*)⟩_**x**,*t*_, as a function of time-interval *τ*. It exhibited oscillatory behavior with a period of about 5 h, indicative of periodic pulsatile movements occurring in the tissue. By calculating the spatial correlation function, ⟨div(**x** + Δ**x**, *t*)div(**x**, *t*)⟩_**x**,*t*_, as function of *r* = |Δ**x**|, we found that the divergence was also correlated in space on a length scale of approximately 200 *µ*m (Figure 1c), which corresponds to approximately 15 cell lengths (see Velocity Field and Correlation Functions section in STAR Methods). Similarly, we noted a spatial decay in the velocity correlation function of ≈ 200 *µ*m that is consistent with the inherent correlation length for divergence (Figure 1f). The velocity field exhibited a temporal decay of a few hours, and is larger than the correlation time of the divergence. This finding is consistent with our observation of regions moving coherently with a relatively uniform velocity on long time scales (Figure 1d, 1e and Figure S3e). Taken together, divergence and velocity correlation plots highlight different collective features of cell motion in the monolayer. Meanwhile we also found that the underlying rotational strength of the flow field was comparable to divergence both in terms of correlation length and time (Figure S3f).

### Pulsatile flows can be modulated by spatially periodic tissue-substrate interaction

We next sought to test whether the period and magnitude of pulsatile flows could be influenced experimentally. Considering the cell monolayer as a layer of active material interacting with the underlying substrate, we reasoned that modulating the tissue-substrate interaction in a spatially periodic fashion would allow to influence the tissue-scale collective flow. To this end, by using micro-contact printing (see Micro-contact printing section in STAR Methods), we deposited the extracellular matrix protein fibronectin as square motifs of controlled dimensions, called *G* − 75 *µ*m, *G* − 150 *µ*m, *G* − 300 *µ*m (see Figures S4a-c, Figures 2a, S5a and S5d, and Movies 3-5). We kept the width of the fibronectin regions *W* = 120 *µ*m to be the same for all three cases while varying the size *G* of the central region. We reasoned that fibronectin coating would locally increase the friction between the tissue and its substrate and separate the entire tissue region into areas of high and low friction (Figure S4a-c). We chose the dimension of these patches *G* + *W* to be around the size of the domains of spontaneous cell pulsations. We then quantified the temporal and spatial correlation of the flow divergence for the entire region. As in the control case, here also we observed the presence of oscillations with period of around 5 h (Figures 2c, S5c, S5f and Figure S4e). But the correlation lengths were less variable across experiments with FN grids, compared to control (Figure S4d), implying that the difference in friction due to FN was tuning the size of pulsatile domains. Remarkably, correlation plots of divergence indicated that the oscillatory component of the flow, as quantified by the dampening of the oscillation in the temporal correlation of the divergence, was maximal when the size of the motifs matched the spontaneous extension of pulsations, in the *G* − 150 *µ*m case (Figure 2, Movie 4). In contrast, when the grid-size was smaller (*G* − 75 *µ*m, Figure S5a-c, and Movie 3) or larger (*G* − 300 *µ*m, Figures S5d-f, and Movie 5), pulsations were less pronounced. It is worth noting that the magnitude of divergence of tissue flow was increased in the larger grids case (≈ 0.1 h^−1^) (Figures 2b and S5e, Figures S5g and S5i) compared with that in smaller grids (≈ 0.05 h^−1^) (Figures S5b and S5h). This observation is consistent with the notion of lower overall friction between the tissue and its substrate when the fraction of substrate area covered by fibronectin is lower.

**Figure 2:**
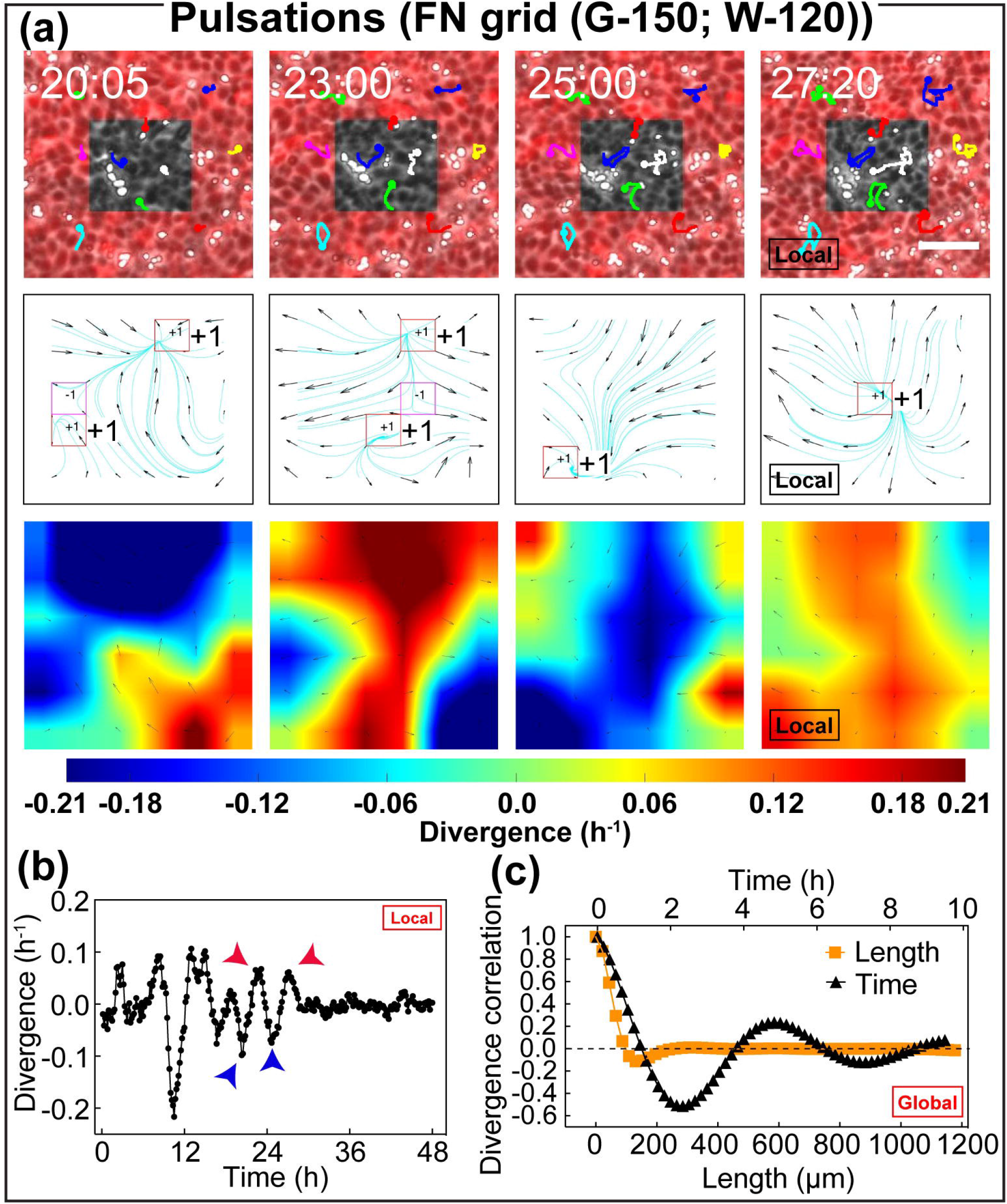
Substrate friction determines the localisation of pulsations. (a)-(b) represent the local characteristics, and (c) illustrates the global characteristics. (a) shows a typical sub-unit of the fibronectin (FN) grid shown in Figure S3b. Here, the first row shows snapshots of one contraction expansion cycle of a pulsatile domain in a single grid unit where tracking highlights the cell trajectories during pulsations. Similarly, the second and third rows show the topological defects and divergence, respectively, for the snapshots in the first row. Color bar indicates the scale for divergence maps where blue and red indicate contraction and expansion, respectively. Scale bar, 100 *µ*m and time in hh:mm. (b) shows the mean divergence plot that highlights the variations in divergence over 48 h for the FN grid domain in (a) and in Movie 4. Arrows indicate data points corresponding to the snapshots in (a). (c) The time correlation function for the corresponding divergence field exhibits striking oscillatory behavior with a period of approximately 5 h. The divergence correlation function in space also shows a minimum at approximately 150 *µ*m, which reflects the periodic pulsatory pattern in space. See also Figures S4, S5 and S9, Movies 3-6.

In these different conditions, more than ≈ 50% of the grids exhibited sustained oscillations. Oscillations lasted for ≈ 48 h, corresponding to a couple of cycles in the *G* − 75 *µ*m case, six cycles in the *G* − 150 *µ*m case, and a couple of cycles in the *G* − 300 *µ*m case. Decay in oscillation amplitude was possibly associated with cell proliferation leading to reduced tissue motion over time. We also noticed that the number of ±1 defects was larger in the *G* − 75 *µ*m and *G* − 300 *µ*m cases than in the *G* − 150 *µ*m case (Figures 2a, Figures S5a and S5d). Finally, in the ‘resonant’ case of *G* −150 *µ*m, we aimed to decrease friction in the central, non-adhesive region of the pattern through incubation with poly(l-lysine) grafted with poly(ethylene glycol) (PLL-g-PEG), which passivates the surface. This process led to ‘agitated’ pulsations, involving localized rapid motions of cells within the passivated square (Movie 6), and this further supports the role of friction in the process. These results strongly suggest that friction can influence the nature of cellular movements. Taken together, these results show that the magnitude of pulsatile flow can be controlled by spatially modulating tissue-substrate interaction.

### Generation of long-range flows in epithelial monolayers

We then examined whether the correlation length of the flow could be experimentally modified. Interestingly, we found that incubation for 12 h with cytoskeleton inhibitors, followed by washing, strongly modulated the correlation length of the tissue-level velocity (see Figures 3a and 3b, Figure S6a-c, and Movies 7 and 8). Specifically, upon the addition of blebbistatin (see Cytoskeletal Inhibitors section in STAR Methods), tissue motion was significantly reduced (Figure S6a). However, about 2 h after subsequent washout of blebbistatin led to the appearance of flows with a longer spatial correlation but without change in persistence over time (Figures 3a-b, Figure S6a, S4h-i; Movie 7), as quantified by the velocity correlation functions. The myosin retrieves its activity within minutes after washout but the cell and the monolayer have their own time to integrate activity and cause flows on 100 *µ*m length scale. A similar effect was also observed following the addition and washout of latrunculin-A (Figures S6b-c, S4h-i and Movie 8). We propose that blebbistatin washout promotes *de novo* monolayer collective order which is distinct from cellular arrangement obtained with standard plating protocol in which cells are randomly distributed at the early time points. We suggest that the formation of new cell-cell contacts after washout resets individual cell polarities to generate long range flow patterns.

**Figure 3:**
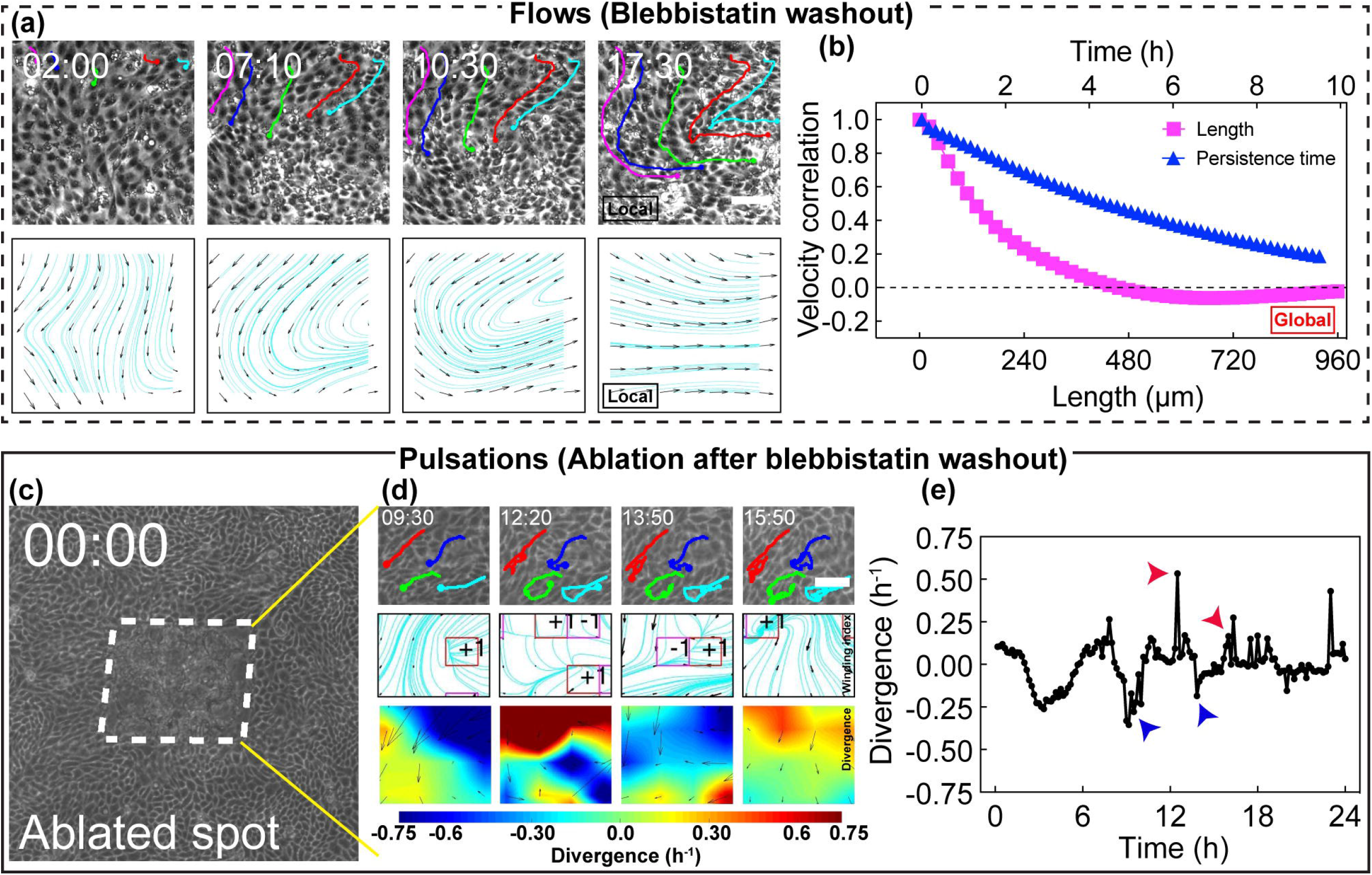
The transition from pulsations to flows and back to pulsations. (a)-(b) Resetting myosin activity (incubation with blebbistatin for 12 h and washout) leads to the transition from pulsations to flows. (a) and (b) highlight the local and global flow characteristics, respectively. The first row of (a) shows snapshots of phase-contrast images where the tracking highlights flow behavior. The second-row shows streamlines without the presence of topological defects. Scale bar, 100 *µ*m. See also the associated Movie 7. (b) shows the spatial and temporal correlation plots for velocity over 36 h. (c)-(e) Ablation of tissue performed after flows that resulted from resetting myosin activity leads to pulsatile behavior. (c) shows snapshot of phase contrast image with the ablated spot highlighted by dotted lines. Scale bar, 100 *µ*m. Time is in hh:mm. See the associated Movie 9 and Figure S6d corresponds to the ablated region in (c). The first row of (d) shows snapshots of phase-contrast images where the tracking highlights back-and-forth pulsatile movements. The second and third rows show winding number and divergence respectively. Scale bar, 50 *µ*m. Time is in hh:mm; (e) shows the mean divergence plot over 24 h for the pulsatile domain shown in (d). Arrows indicate the data points corresponding to snapshots in (d). See also Figures S6 and S9, Movies 7-9.

We had observed that pulsations in the MDCK monolayer started around the time when the cells were about to become confluent by the closure of gaps or ‘wounds’ in the tissue. Assuming that an empty space can lead to pulsatile motion of cells, we generated a wound using laser ablation after washout of the cytoskeletal inhibitor when the tissue exhibited a strongly coherent flow (see Figure 3c, Figure S6d, and Optical setups and imaging conditions section in STAR Methods). Interestingly, cells then underwent back and forth motions with contraction and relaxation after ablation, in a way reminiscent of pulsatile flows observed without treatment (Figure 3d and Movie 9). This observation was further confirmed by the measurement of divergence and its correlation (Figures S4f and S4g, Figure 3e and Figure S6e). These results indicate that a combination of cytoskeletal remodeling and physical perturbations can toggle collective cell movements between pulsatile and long-range flows.

### Cellular mechanisms behind pulsatile flows

Following the same idea as laser ablation, where wound closure led to pulsations, we then sought to generate controlled wounds in the tissue to test their effect on the flow properties. Cells were plated on the substrate with microfabricated pillars [34], which were then removed, generating regularly placed empty spaces in the tissue. We then noticed that cells around the site of wound closure underwent pulsatile flows (Figures S7a and S7b, and Movie 10). We took advantage of this setup to visualize the cellular myosin distribution in cells participating in pulsatile flows (Figure 4a). We noticed that contracting zones appeared to exhibit an increased density of myosin. We also observed the transient appearance of a radial gradient of myosin fluorescence intensity around the wound, with larger intensity values towards the center of the wound in the contracting phase. This gradient was more homogeneous in the expanding phase (Figures 4b and 4c, Figures S7c and S7d, and Movie 10). These results suggest that myosin gradients participate in driving pulsatile flows in the tissue.

**Figure 4:**
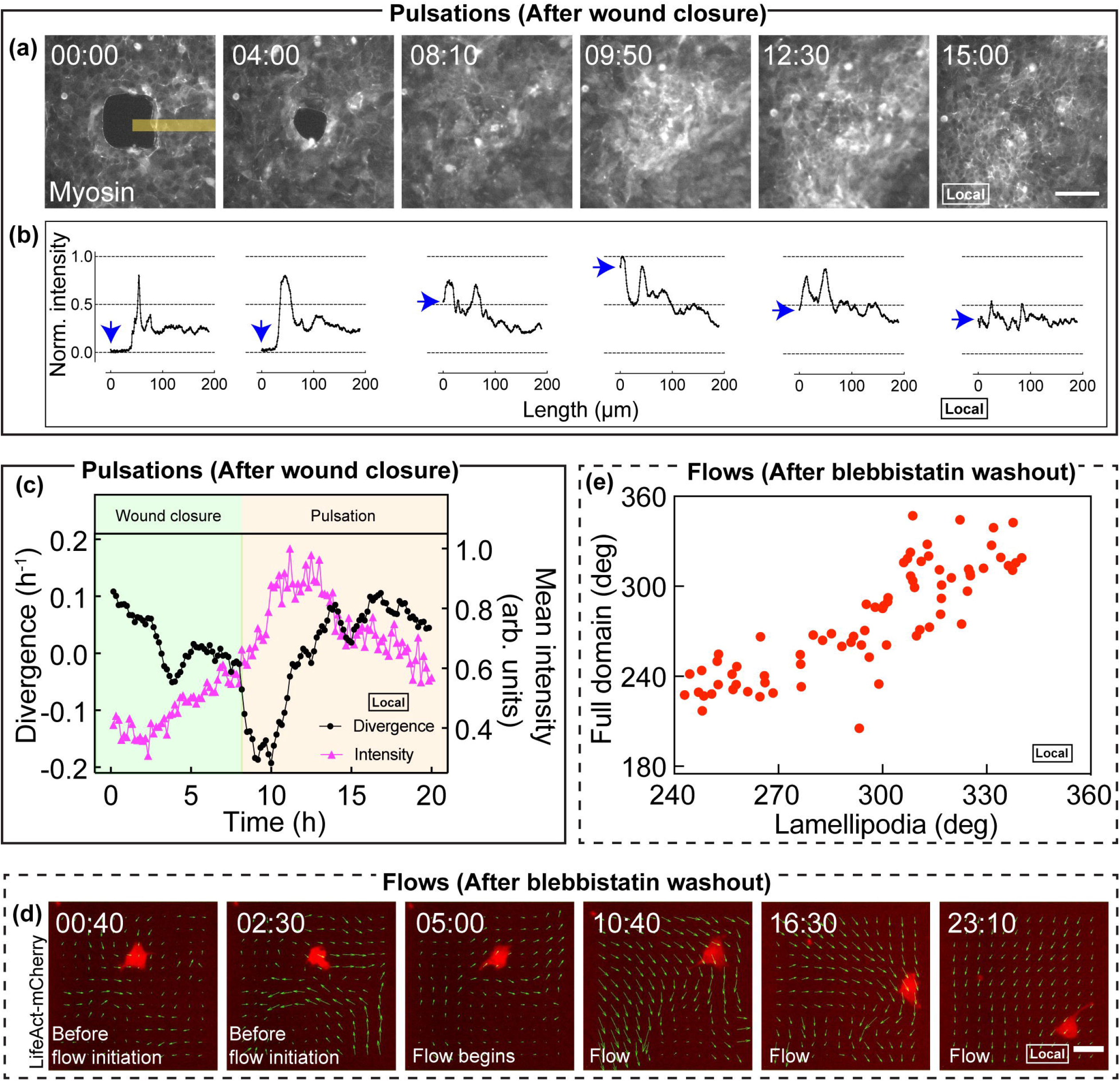
Molecular actors associated with pulsations and flows. (a)-(c) Fluctuation in myosin density during the contraction and expansion phases of pulsation. Micropillar assay was used to create wounds in the monolayer. Following wound closure, pulsations were observed in these sites. (a) Snapshots showing the fluctuation in myosin-GFP intensity. The first and the second time points show the wound and its eventual closure, respectively. Subsequent time points show the densification and diffusion of myosin. Scale bar, 100 *µ*m and time is in hh:mm. (b) shows myosin intensity profiles obtained along the yellow stripe in (a). Myosin intensity is initially zero at the center (0 *µ*m) of the wound (indicated by blue arrows) and increases to the half-way mark during wound closure. However, as the pulsation begins with contraction (at 12:40 in (a)), myosin intensity reaches the maximum and diffuses back to the half-way mark as the domain extends. See the associated Movie 10, Figure S7a and Figure S7b. (c) Divergence and intensity plots for the pulsatile domain shown in (a). The green region represents the wound closure phase, and the pink region represents the pulsation phase. (d)-(e) Orientation of lamellipodia aligning with the flow direction (on resetting myosin activity through incubation with blebbistatin for 12 h and washout). The first two snapshots in (d) represent the timepoint before flow initiation; the transition to flows is shown by the third time point and the subsequent time points show the flow phase. (d) The lamellipodial orientation of a cell transiently transfected with LifeAct-mCherry, shown in red, is superimposed with the direction of flow field vectors (green). Scale bar, 50 *µ*m, and time is in hh:mm. (e) Plot showing the linear tendency between lamellipodia orientation and the mean flow direction of the domain. The lamellipodia direction (obtained from the effective cell direction - see Velocity-polarisation correlation for migrating cells section in the STAR methods) in the *x*-axis, is obtained for the cell shown in (d). See also Figures S7 and Movies 10 and 11.

We also tested whether cell lamellipodia orientation was correlated with tissue flow. To follow lamellipodia dynamics, we used mosaic experiments with cells transiently transfected with Lifeact-mCherry (Figures 4d, Figure S7e, and Movie 11; also see Cell culture and sample preparation section in STAR Methods). Using blebbistatin washout to induce large-scale correlated flows, we found that lamellipodia direction indeed correlated with the movement of cells (Figures S7e and S7f, Figures 4e, S7g and S7h, see Velocity-polarisation correlation for migrating cells section in STAR Methods). We chose to quantify the lamellipodium orientation by tracking and segmenting the outline of cells labelled with Lifeact-mCherry and calculated a nematic order parameter from the cell contour. We found that the nematic axis associated with cell elongation was then typically aligned with the direction of the flow (see Velocity-polarisation correlation for migrating cells section in STAR Methods, Figure S2). These observations support that lamellipodia extend along the direction of flows.

In summary, these observations support the idea that pulsatile contraction and expansion are associated with variations in cell myosin intensity and that cells are polarized along the direction of the flow.

### Numerical simulations to model tissue pulsatile flows

To identify the origins of the pulsatile collective flow, we ran numerical simulations of the tissue flow based on a vertex model [35]. In our model, cells were assigned a polarity vector that evolved over time according to a local alignment rule and a polarity diffusion. The cells were subjected to a motile force along the polarity axis (Figure 5a).

**Figure 5:**
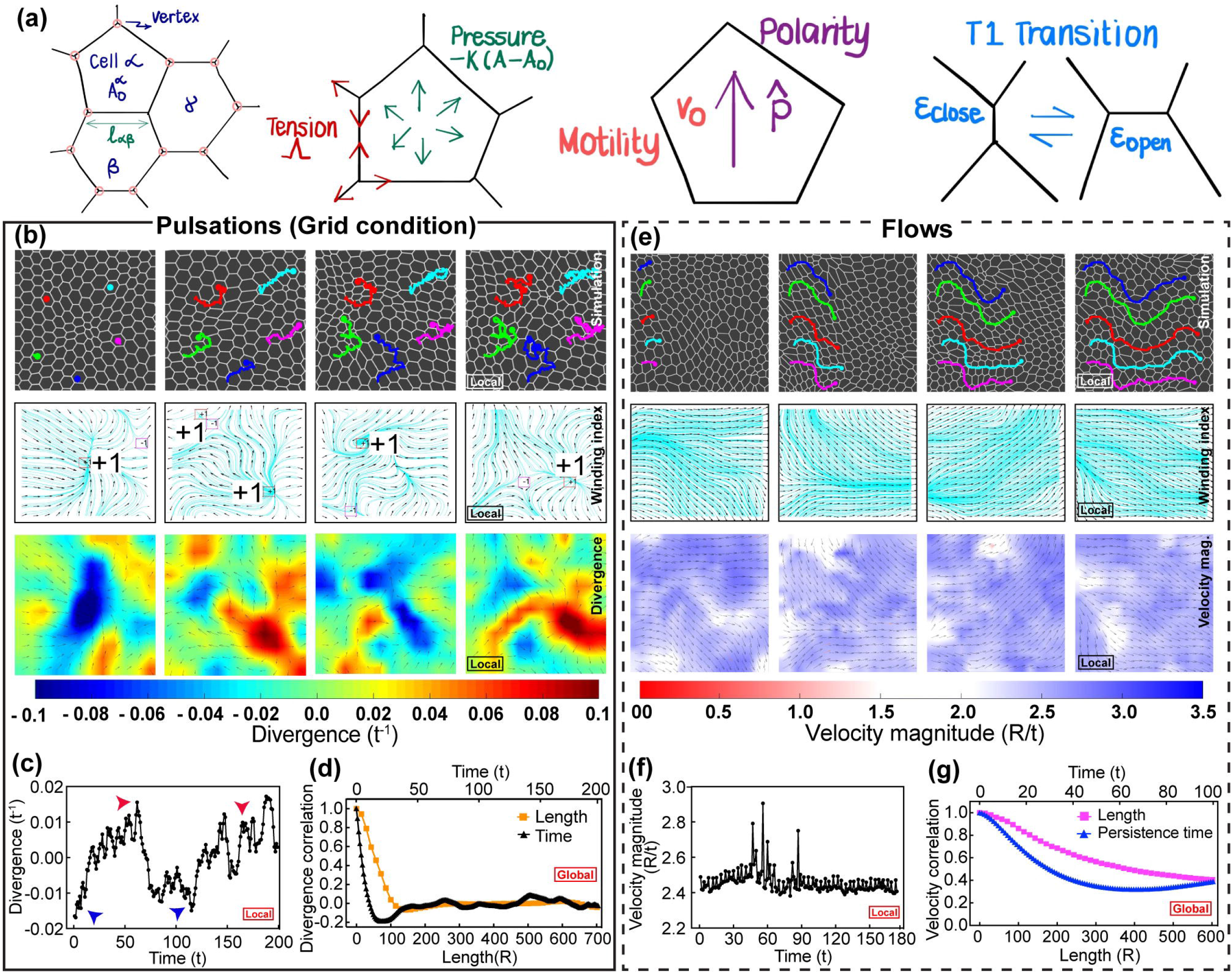
Simulations of pulsations in friction grid and flows using active vertex model. (a) Basic schematic of the vertex model and the source of forces on the vertices. The outcome of simulations for friction grid (b)-(d) and flows (e)-(g) correspond to their experimental counterparts in Figures 2 and 3, respectively. In (b), the first row shows: periodic contraction and expansion of a local domain with the back-and-forth movement of cells. The second and third rows show the topological defects from winding number analysis and divergence, respectively. Color coding: blue indicates contraction and red indicates expansion in the panels showing divergence field. See the associated Movie 13. (c) Total divergence for the local domain shown in (b) as a function of time. (d) The time and space correlations for divergence over the entire simulation. As observed in the experimental findings, length and time scales emerged from the divergence correlation plots. In (e), the first row shows the presence of long-range flows in the selected region. The second and third rows show the absence of topological defects and velocity magnitude, respectively. Color coding: red and blue indicate low and high velocities in those panels representing velocity magnitude. See the associated Movie 15. (f) The corresponding velocity magnitude within the domain in (e) is nearly constant in time. (g) Velocity correlation function over the entire simulation shows persistence (long correlation) in both space and time. For a detailed discussion on experimental units of time and length, see Additional Details for the Computational Mode section in the STAR Methods. See also Figure S8 and Movies 12-15.

In the basic vertex model of the confluent monolayer, the epithelial cells were represented as polygons that were created with vertices (Figure 5a). Every pair of adjacent vertices was shared by an edge which in turn had a cell on each of its side. The position of the vertices and the connectivity of the cells defined the geometry and topology, respectively, of the tissue. The mechanical properties were assigned to the minimal vertex model via a work function *W* :

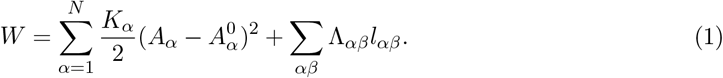

The first term of the work function modeled the isotropic mechanical resistance with stiffness *K*_*α*_ from a cell *α* having actual area *A*_*α*_ to any deviations from its preferred area 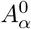. The second term represented the effective interaction energy between two connected cells *α* and *β* that resulted from a combination of cell-cell adhesion energy (Λ_*αβ*_) and acto-myosin contractility along the shared edge *αβ* length *l*_*αβ*_. The force acting on a particular vertex *i* with position **r**_*i*_ was

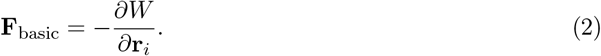

The cells within the colony were polarised and had a tendency to migrate (Figure 5a). The motility behavior of an isolated polarised cell was modeled using a simple description in which the cell moved with speed *v*_0_ along its polarity direction 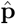. For the vertex model, this self-propulsion tendency of cells was converted to an effective motile force on its constituent vertex *i* as

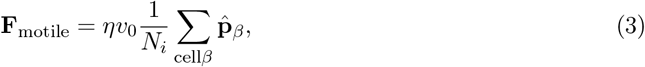

where summation was over the polarities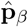, respectively, of the *N*_*i*_ cells that contain the vertex *i*, and *η* is the effective friction coefficient between the cell vertex and the substrate [35, 36]. Initially, *N* = 3060 cells were created in a periodic box of size *L*_*x*_ × *L*_*y*_. Each cell is provided a polarity 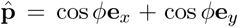, where the angle *ϕ* was chosen randomly from [0, 2*π*). The time evolution of vertex positions **r**_*i*_ was based on the equation

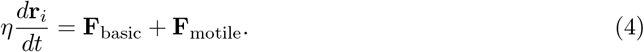

To temporally evolve the polarity of the cells we utilised the experimental observations depicted in Figure S2. Specifically, we assumed that the polarity vector 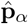for a cell *α* has two tendencies – (i) to align with the direction 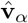of its instantaneous velocity and (ii) to undergo rotational diffusion – which mathematically translated to the following differential equation [37, 38]

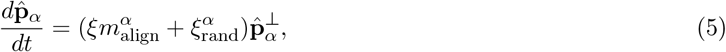

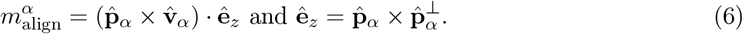

Here, *ξ* was the alignment strength of polarity and *ξ*_rand_ was the diffusive, rotational, Gaussian noise with strength *D*_*r*_. The differential equations were solved using an explicit forward scheme with a time-step Δ*t*. See STAR Methods for additional details. Our model had seven parameters: preferred cell area (*A*_0_), cell area modulus (*K*_0_), line tension of cell interfaces (Λ), cell motility (*v*_0_), polarity alignment rate (*ξ*), rotational diffusion of polarity (*D*_*r*_), and substrate friction coefficient (*η*). From these parameters, a set of time scales can be defined (see STAR Methods in the Additional details for the computational model section).

We first tested the simulations with periodic boundary conditions and no rotational noise on the cell polarity vector and failed to generate pulsations. However, flows of cells were observed in many of the versions [39]. This finding suggests that the generation of pulsations required another element, provided here by the rotational noise on cell polarity. We found that the absolute and the relative values of *ξ* and *D*_*r*_ were important in setting the nature of collective migration modes in the tissue (see Additional details for the Computational Model section in STAR Methods). High values of *ξ* promote polarity alignment in some direction, and hence long-range flows. On the contrary, high values of *D*_*r*_ promoted fast and short-range oscillations. When the relative values of *D*_*r*_ and *ξ* were well-balanced, we found temporally periodic and spatially oscillatory migration patterns of cells similar to that experimentally observed.

We next sought to reproduce three experimental conditions (Figure 5, Figure S8, and Movies 12-15). We numerically mimicked standard monolayer (Figure S8a-c, Movie 12), monolayer on grid (Figure 5b-d and Movie 13), and flows after blebbistatin washout (Figure 5e-g and Movie 15). To perform these numerical experiments, we tested the parameters that could allow reproduction of the experimental data. To be quantitative, we also plotted the divergence correlation and the velocity correlation as a function of time and space. The three situations are represented in Figure 5d, Figure 5g, and Figure S8c, all of which show good agreement with our data. We needed to have 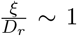, grid-dependent friction, and 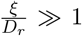to produce, respectively, Figures S8b, S8f and S8c, Figures 5c, S8d and 5d, and Figures 5f, S8e and 5g. Interestingly, we could also mimic the agitated movement of cells observed in a fibronectin grid with a PEG coating at the center, which is expected to reduce cell-substrate friction in that region (Movie 14). We note that it is the differential friction between the central region and the adhesive patch of the simulated grid that leads to the modification in the pattern of cellular movements. Interestingly, the velocity patterns were more pronounced when the difference in the friction between the two regions was greater. We use periodic boundary conditions for all the simulations. In Movies 12-15 in which simulations are compared with their experimental counterparts, a smaller spatial domain is extracted from the simulation region of the complete tissue.

Our model assumptions and predictions are consistent with experimental observations. When the velocity direction and polarity at any position in monolayer were weakly correlated, pulsation was the main mode of cell movements (Figures S9a-S9b and Velocity-polarisation correlation for migrating cell section in the STAR Methods). In contrast, when the velocity-polarity correlation was strong, cell flows were dominant (Figures S9c-S9d). In our model, the velocity alignment term *ξ* tends to align cell polarity with its velocity direction, whereas the rotational noise *D*_*r*_ perturbs this alignment. Taken together, these observations support our modeling results that when velocity alignment strength is dominant (*ξ/D*_*r*_ ≪ 1), we obtain long range cellular flows. However, when the strength of velocity alignment and random rotation are comparable (*ξ/D*_*r*_ ∼ 1), we generate pulsatory cell movements.

Taken together, these results reproduce the main observations of our experiments. We conclude that a simple combination of cell motility and polarity dynamics could account for the pulsatile nature of the flow. More specifically, modulating substrate friction, polarity diffusion and the strength of polarity-velocity alignment led to a variations in tissue kinematics.

## Discussion

Pulsations and flows in tissues are striking examples of collective motions. Although they have independently been studied earlier in MDCK monolayers [26, 27, 40–46], in this paper, we generated both types of multi-cellular dynamics in the same epithelial monolayers. We controlled each of them by spatially modulating substrate friction and by introducing in and then washing-out cytoskeletal inhibitors. We revealed that these perturbations affected the external environment and internal cellular properties of the monolayers, leading to these distinct robust collective motions. We also successfully reproduced these dynamics in a minimal vertex model with cell motility that took rotational diffusion of polarity and polarity-velocity alignment as essential components. In this model, a simple change in respective weights of these two polarity mechanisms resulted in a switch between pulsations and long-range flows, showing a way to control collective cell motions in monolayers. Thus, our study shows that the very same epithelial layer can switch its dynamics using simple rules, and pulsations and flows appear as emergent properties of epithelial monolayers. Analogously, we could expect that tissues *in vivo* could also switch their dynamics in time and space with the novel mechanisms we report by tuning local friction or modulating polarity dynamics in the same organ. Future studies can test these hypotheses *in vivo* by tuning the local interaction between the neighbouring tissues by locally overexpressing ECM proteins or by locally incubating organs with myosin inhibitors and visualizing their recovery live. Thus, our findings open up a new perspective on using these readouts and numerical simulations to test morphogenetic events in general.

It is interesting to note that the domains for pulsations and flows are about 100 *µ*m in length scales for time scales of about few hours. These similar values can be obtained by scaling arguments. The Maxwell time *τ* is the ratio of viscosity over Young’s modulus. Taking viscosity of about 10^7^ Pa.s and Young modulus of about 10^3^ Pa [47, 48], we obtain *τ* of the order of hours. Cell motile speeds are about *v*_0_ ∼ 20 *µ*m*/*h. The product *τ* × *ν* _0_ gives *l* ≈ 100 *µ*m. Altogether, these estimates support a simple viscoelastic behavior.

Collective cell migrations *in vivo* occur in an integrated manner, involving signaling and mechanics pathways. For example, in zebrafish, that the so-called lateral line undergoes collective migration through chemical gradients generated by the tissue itself [4, 49]. Similarly, in *Xenopus laevis* neural crest cells, stiffening is involved in setting the mesenchymal to epithelial transition through mechanisms involving an interplay between tissue mechanics and local signalling [50, 51]. Still, mechanical events alone were sufficient to capture some morphogenetic events. Hence, the search for physical rules is essential to capture the deformation kinematics of tissues. For instance, cell motility per se was associated with the elongation of the amniote axis. Similarly, differential random motion of cells was reported to control somitogenesis in chicken embryos [52–54]. Even if gradients in fibroblast growth factor (FGF) signaling were also present as well in this case, the cellular dynamics were sufficient as the main driving force for elongation, supporting the need to look for physical principles piloting morphogenesis. Germ cells were also shown in many organisms to move towards the gonads through active and passive processes associated with the dynamics of the surrounding tissues [55, 56]. Our generic mechanisms of tissue dynamics for identical cell lines could serve as a framework to revisit these examples guided by the quantification procedure and theoretical methods developed in this study.

In this context, we herein report two types of collective cell motion and provide a generic framework to explain them. Pulsatile movements were observed in various conditions. For example, in *Drosophila*, pulsed contractions guided by acto-myosin are involved in apical constriction and abdominal morphogenesis [12, 57, 58]. Moreover, oscillation may vary across the tissues, as shown in *Drosophila* abdominal morphogenesis [58], and the accompanying myosin gradients are reminiscent of the ones we report in the current study. Elsewhere, oscillations within filopodia during the heart formation of *Drosophila* are essential to optimize morphogenesis in a proof-reading mechanism [59, 60]. Cardiomyocytes also undergo these oscillatory motions both *in vivo* and *in vitro* [61]. This expanding list of pulsation incidences suggests that a self-organized ‘pacemaker’ associated with a simple pulsation mechanism of physical origin, which span various tissues, model organisms, timescales, and length scales, could be involved in these situations.

We propose the same extension from our study for describing flows *in vivo* [62]. A previous study has shown how forces drive epithelial spreading in zebrafish gastrulation through a flow-friction mechanism [7]. During epiboly in the insect *Tribolium castaneum*, part of the tissue becomes solid and pulls the associated tissues undergoing rearrangement and direct flow [8, 63]. A similar mechanism was suggested during wing formation in *Drosophila* with the hinge and the connected blade [6]. Force balance associated with adhesion between the blastoderm and the vitelline envelope led to this long-range flow. In another context, acto-myosin cables were shown to lead to collective movement of cells in a manner reminiscent to a liquid flow [64]. These situations show similar cell flows to that in the present study. Our study suggests that a common mechanism across scales, tissues, and species may be related to the greater contribution of velocity alignment than rotational diffusion in polarity. Lamellipodia in cells and its actin dynamics could be the main motor in these long-range flows.

This search for simplicity in the multi-cellular mode aims at extracting generic principles of morphogenesis. The MDCK epithelial monolayer in this study is an excellent system to mimic any epithelial dynamics *in vivo*. As mentioned above, collective cellular motions and deformations are common in developmental biology. However, the focus is generally on the genetic and signaling pathways associated with their onsets. Our *in vitro* results suggest that our readouts of diffusion in polarity and cell alignment can be used *in vivo* to test mechanisms in complex systems during morphogenesis. Because these parameters are now identified, collective cell dynamics can be tested with the same framework across species. Considering the large number of model systems available, including *Drosophila, C. elegans*, to zebrafish, *Xenopus*, and mice, these self-organization rules could be explored experimentally and compared across organs and species.

Our current study focused mainly on the physical parameters (substrate friction, cell motility, diffusion in polarity, and polarity-velocity alignment) controlling these types of motion and their cellular readouts through lamellipodia distributions and myosin gradients. In the future, it would be interesting to study the coupling between mechanical processes and cellular signaling, with experimental measurements of both types of events and their inclusions in the associated computational framework [65]. Indeed, recent works have highlighted the connection of wave propagation in tissues with signaling. Mechanical events alter signaling activity and these feedbacks are mediated by mechanochemical interactions between cells through extracellular-signal-regulated kinase (ERK) activation [65]. Investigations on such interplay between mechanical and chemical contractions and waves could generate new paradigms in morphogenesis and allow the identification of conserved principles of mechanics and signaling networks in living matter. Cell density is another important parameter that regulates the characteristics of monolayer dynamics in a time dependent manner [21]. In our experiments, beyond 48 h, cell proliferation leads to jamming like effect where the large scale dynamics are arrested and only local movements are observed. Along this line, the role of cell density in transitioning between pulsations and flows is an interesting avenue to be explored both in terms of dynamics [21] and the underlying molecular mechanisms [66].

### Limitations of the study

In the current study, we generate cellular flows by washing out the myosin inhibitor blebbistatin of the confluent monolayer but do not perfom experiments to directly test the associated mechanisms. Future experiments could modulate cell-cell interactions by incubating DECMA-1 antibody to perturb cell-cell interactions after blebbistatin washout. We anticipate that at high concentrations of DECMA-1, cell-cell junctions would be separated whereby no collective motion would be observed. However, at intermediate detachment levels, we expect that the spatial length and persistent time of the cellular flows would be decreased. In contrast, other MDCK cell lines overexpressing cadherin could be generated in which case we expect that both the flow correlation length and persistent time of the flows would be larger. These experiments would help substantiate the key role of cell-cell junction lifetime in modulating the velocity alignment of cells.

Our experimental findings in this paper are based on MDCK epithelial monolayers. However, for a more general understanding, other epithelial cells such as HaCat and Huvec cells could be used. Nevertheless, based on our understanding from MDCK experiments and simulations, we predict that the interplay between the levels of adhesion (cadherin) and force (myosin) would determine the types of dynamics – pulsations or waves. We anticipate the cells with higher levels of adhesion to facilitate polarity alignment and cellular flows, whereas cell lines with the larger forces may facilitate pulsations. At that stage, numerical simulations would allow to quantitatively test these predictions by associating appropriate model parameters.

### Future work

In the current study, we do not systematically explore the role of myosin levels on tissue kinematics. For that, clones with controlled levels of myosin could be prepared in future experiments. The role of myosin levels on the length and time-scales of cellular flows after blebbistatin washout experiment could also be explored from these experiments. We propose that the over-expression of myosin would decrease the persistence length and time by reducing velocity alignment whereas downregulation of myosin would trigger the opposite results. Future experiments would also help disentangle the potential interplay of myosin roles in velocity alignment and polarity diffusion as is proposed theoretically.

In this work, we do not perform a detailed comparative study of the combined quantitative effect of myosin and cadherin on collective cell movements (pulsations or flows) observed in the monolayer. Mosaic experiments using the clones discussed above would allow to further probe the respective mechanisms that are involved. By using fluorescent reporters for cadherin and myosin with distinct colours, levels of adhesion and force could be measured with respect to the local patterns of monolayer kinematics. From such measurements, a more precise mapping between the pattern of collective cell migration and spatiotemporal levels of myosin and cadherin can be obtained.

In the current work, we do not explore the role of substrate stiffness in quantitatively dictating the patterns of collective cell migration. However, we propose predictions about how the cell migration patterns would be affected by the modulation of substrate stiffnesses. With soft surfaces, we expect pulsations to be facilitated based on the fact that deformations of substrates can accompany the contractile activity of cells. In this case, the oscillation period and the spatial length could both be larger. Analogously, we expect a similar increase in the length and time scale for flow conditions. With stiffer substrate, however, we do not expect major changes since our substrates are already much above the typical Young modulus of our epithelial monolayers.

## Supporting information

Resource table

## Author contributions

R.T. designed and performed experiments, analyzed data, and wrote the manuscript. A.B. performed experiments. G. S. designed theoretical model, analyzed data, and wrote the paper. M.M.I. designed theoretical model, performed simulations, analyzed data, and wrote the paper. D.R. designed the study and experiments, supervised the project, analyzed data, and wrote the paper.

## Acknowledgements

We thank Frank Jülicher, S. S. Soumya, and Vikram Gadre for helpful discussions, and Romain Levayer and Julien Vermot for important feedbacks on morphogenesis in developmental biology. We thank the Riveline Lab for discussions and help. We acknowledge the Imaging Platform from IGBMC and P. Reilly for critical reading of the manuscript. We are grateful to W. James Nelson, Shigenobu Yonemura and Roland Wedlich-Söldner labs for sending cell lines. D.R. acknowledges support from CNRS (ATIP), ciFRC Strasbourg, the University of Strasbourg, Labex IGBMC. This study with the reference ANR-10-LABX-0030-INRT has been also supported by a French state fund through the Agence Nationale de la Recherche under the frame programme Investissements d’Avenir labelled ANR-10-IDEX-0002-02. R.T. was an IGBMC International PhD Programme fellow supported by LabEx INRT funds. M.M.I. acknowledges funding from Science and Engineering Research Board (MTR/2020/000605). G.S. was supported by the Francis Crick Institute which receives its core funding from Cancer Research UK (FC001317), the UK Medical Research Council (FC001317), and the Wellcome Trust (FC001317). D.R. and M.M.I. also acknowledge hospitality at MPI-PKS, Dresden, where a part of this work was done.

## Main figure titles and legends

## Movie captions

**Movie 1:**. Pulsatile domain extracted from the MDCK-E-cadherin-GFP monolayer, related to Figure 1. Cells spontaneously exhibit collective contraction and extension phases upon reaching confluency. Time in hh:mm.

**Movie 2:**. Domain of cells exhibiting flows, extracted from MDCK-E-cadherin-GFP monolayer, related to Figure 1. Cells display mainly flow behaviour. Time in hh:mm.

**Movie 3:**. FN grid (G-75; W-120) condition, related to Figure 2. Pulsatile domain extracted from MDCK-E-cadherin-GFP monolayer plated on fibronectin grids of width (W) 120 *μ*m and a gap (G) (non-fibronectin area) of 75 *μ*m. Fibronectin grid is shown in red and the gap is shown in grey. Time in hh:mm.

**Movie 4:**. FN grid (G-150; W-120) condition, related to Figure 2. Pulsatile domain extracted from MDCK-E-cadherin-GFP monolayer plated on fibronectin grids of width (W) 120 *μ*m and a gap (G) (non-fibronectin area) of 150 *μ*m. Fibronectin grid is shown in red and the gap is shown in grey. Time in hh:mm.

**Movie 5:**. FN grid (G-300; W-120) condition, related to Figure 2. Pulsatile domain extracted from MDCK-E-cadherin-GFP monolayer plated on fibronectin grids of width (W) 120 *μ*m and a gap (G) (non-fibronectin area) of 300 *μ*m. Time in hh:mm.

**Movie 6:**. FN grid (G-150; W-120) condition with passivation, related to Figure 2. Pulsatile domain extracted from MDCK-E-cadherin-GFP monolayer plated on fibronectin grids of width (W) 120 *μ*m and passivated regions (G) (pLL-g-PEG coating) of 150 *μ*m. White box shows the passivated area. Time in hh:mm.

**Movie 7:**. Blebbistatin washout condition, related to Figure 3. The movie initially shows a pulsating domain in control condition from MDCK-E-cadherin-GFP monolayer. After addition of 100 *μ*M blebbistatin, pulsations are arrested. When blebbistatin is washed out after 12 h incubation, the same domain starts to exhibit long range flows. Time in hh:mm.

**Movie 8:**. Latrunculin-A washout condition, related to Figure 3 and S6. The movie initially shows a pulsating domain in control condition from MDCK-E-cadherin-GFP monolayer. After addition of 1 *μ*M latrunculin-A, pulsations are arrested. When latrunculin-A is washed out after 12 h incubation, the same domain starts to exhibit long range flows. Time in hh:mm.

**Movie 9:**. Transition from flows to pulsations, related to Figure 3. After 12 h after blebbistatin washout, a large wound (≈ 150 × 150 *μ*m^2^) is induced by laser ablation in MDCK-E-cadherin-GFP monolayer. The dotted box in the beginning of movie shows the ablated spot. During the wound healing process that followed laser ablation, cells exhibited oscillatory phase highlighted by smaller continuous rectangle in the top right. Time in hh:mm.

**Movie 10:**. Myosin density fluctuation during pulsation, related to Figure 4. Pulsatile domain extracted from MDCK-myosin-GFP monolayer. Monolayer undergoes wound closure with removal of micro-pillars followed by pulsatile activity. Phase contrast images of the cells are shown (left) and GFP-tagged myosin (right). During the pulsation that follows wound closure, accumulation of myosin can be seen with contraction. The accumulated myosin diffuses back to original state during extension phase. Time in hh:mm.

**Movie 11:**. Orientation of lamellipodia with the flow field, related to Figure 4. Domain exhibiting flow behaviour extracted from MDCK-E-cadherin-GFP monolayer transiently transfected with LifeAct-mCherry. The cell shown in red is transfected with LifeAct-mCherry. The flow field vectors are shown in green. Time in hh:mm.

**Movie 12:**. Isolated pulsatile domains from the simulation (right) and experiment (left), related to Figures 5 and 1. The parameters for the simulation are: *ξ* = 0.05, *D*_*r*_ = 0.02, *v*_0_ = 0.1, *μ* = 1*/η* = 0.001, *ϵ*_close_ = 2, *ϵ*_open_ = 3, *K* = 0.003, Λ = 100.

**Movie 13:**. Isolated domains from simulation (right) and experiment (left) for the FN grid condition, related to Figures 5 and 2. The FN grid condition is simulated by increasing the motility (*v*_0_) and mobility (*μ* = 1*/η*) of the cells in the center relative to those at the periphery (FN coated). The parameters for the simulation are: 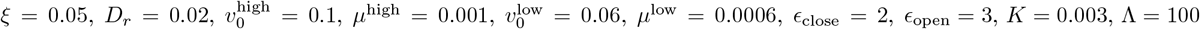. The dimension of the central square and the total thickness (left + right) of the fibronectin grid is kept to be the same.

**Movie 14:**. Isolated domains from simulation (right) and experiment (left) for the FN grid condition with PEG, related to Figures 5 and 2. The FN grid condition is simulated by increasing the motility (*v*_0_) and mobility (*μ* = 1*/η*) of the cells in the center relative to those at the periphery (FN coated). The contrast in these properties between the central and the peripheral region is enhanced when compared to the regular grid in Movie 13. The parameters for the simulation are: 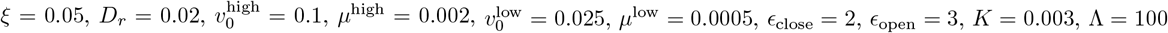. The dimension of the central square and the total thickness (left + right) of the FN grid is kept to be the same.

**Movie 15:**. Isolated domains from simulation (right) and experiment (left) for the flows, related to Figures 5 and 3. The parameters for the simulation are: *ξ* = 0.05, *D*_*r*_ = 0.003, *v*_0_ = 0.1, *μ* = 1*/η* = 0.001, *K* = 0.003, Λ = 100.

## RESOURCE AVAILABILITY

### Lead contact

Further information and requests for resources and reagents should be directed to and will be fulfilled by the lead contact, Daniel Riveline.

### Materials availability

This study did not generate new unique reagents.

### Data and code availability

- All data generated or analysed in this study are available from the lead author upon reasonable request.
- Codes to analyse the data and perform numeric calculations are available from the lead author upon reasonable request.

## EXPERIMENTAL MODELS AND SUBJECT DETAILS

### Cell culture and sample preparation

MDCK cells labelled with Green Fluorescent Protein (GFP) for E-Cadherin (gift from Nelson lab, Stanford University), MDCK cells labelled with actin-mcherry/myosin-GFP (gift from Roland Soldner lab), and MDCK cells labelled with GFP for myosin (gift from Shigenobu Yonemura lab, RIKEN) were cultured at 37°C in low-glucose Dulbecco’s modified Eagle’s medium (DMEM) supplemented with 10% Fetal Bovine Serum (FBS) and 1% Penicillin-Streptomycin antibiotics. Cells were maintained at a minimal confluency of 40% − 50% and then trypsinized at 70% − 80% confluency for experiments. Live experiments were performed in Leibovitz L-15 medium and the initial seeding density was maintained at 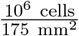 for all experiments. For the mosaic experiment, MDCK-Ecadherin-GFP cells transiently transfected with LifeAct-mCherry were mixed with non-transfected cells.

## METHOD DETAILS

### Cytoskeletal Inhibitors

To inhibit cytoskeletal proteins, these inhibitors were used at the indicated concentrations: blebbistatin (100 *μ*M) and latrunculin A (1 *μ*M). For incubation with inhibitors, media from the dish was replaced with media with the inhibitor. During washout, inhibitor media was carefully removed and the sample was rinsed at least twice with fresh L-15 before finally adding fresh L-15 for further acquisition. In both blebbistatin and latrunculin A conditions, the inhibitor was added to the monolayer after 14 h - 16 h of reaching confluency. The monolayer was then incubated with the inhibitors for 12 h and then washed out. Flows were observed in the following ≈ 36 h starting ≈ 2 h after washout.

### Micro-contact printing

Micropatterning of fibronectin on glass coverslips was performed using the micro-contact printing procedure [67]. First, motifs were microfabricated as SU-8 molds on silicon wafers using standard photolithography technique. Next, the replica were molded on polydimethylsiloxane (PDMS) at a pre-polymer, crosslinker ratio – 9 : 1 (V/V)) for micropatterning. During the micro-contact printing procedure, coverslips were hydrophilized with “Piranha” activation [Piranha: 3:7 parts of H_2_O_2_ and H_2_SO_4_ followed by functionalization with (3-mercaptopropyl)trimethoxysilane for 1 h by vapor phase deposition. The coverslips were then dried at 65°C for 2 h. Next, PDMS stamps with microstructures were hydrophilized by oxygen plasma activation (Diener Electronic) and incubated with TRITC or HiLyte 488-labelled Fibronectin (10 *μ*g*/*ml) for 1 h. The dried stamp was then brought in contact with the activated coverslip and a gentle pressure was applied to transfer the fibronectin patterns from the PDMS stamps to the coverslip. The coverslip was then stored in phosphate buffered saline (PBS) solution at 4°C until the time of cell seeding. If the non-printed zones were to be passivated, the micropatterned coverslip was incubated with PLL-g-PEG (poly-L-Lysine-grafted-PolyEthylene Glycol) (100 *μ*g*/*ml) in 10mM HEPES (pH 7.4) solution for 20 min. Then the coverslips were subsequently rinsed with PBS before proceeding for cell seeding.

### Wound healing experiments

Wound healing experiments were performed using microfabricated pillars [34]. A PDMS stamp with micropatterns was incubated with 0.2% pluronic acid for 1 h and then rinsed with PBS. The stamp was then activated using an O_2_ plasma cleaner and the side with micropillars was brought in contact with an activated glass coverslip for bonding. The bonded stamp and coverslip were stored at 65°C for 30 min to strengthen the bonding. After UV sterilization of the stamp and coverslip, cells at a concentration of approximately 10^6^ cells*/*20 *μ*l were added at the interface between the stamp and the coverslip. The cells filled the space between the micropillars and they were allowed to settle for 45 min. Once the cells have settled, fresh DMEM media was added to the sample which was then stored at 37°C for ≈ 12 h to allow cell spreading between the micropillars. After ≈ 12 h, the PDMS stamp was carefully peeled off from the coverslip and the sample was finally rinsed and filled with L-15 media before imaging was started.

### Optical setups and imaging conditions

Acquisition by phase contrast and fluorescence microscopy was performed using inverted Olympus CKX41 microscopes. The setups were equipped with a manual stage or a Marzhauser Wetzler stage with a stepper motor (MW Tango) enabling multi-point acquisitions, and an Xcite Metal-halide lamp or OSRAM mercury arc lamp for fluorescence acquisition. The phototoxicity during acquisition was prevented by using two shutters (Uniblitz VCM-D1 / VMM-D1 or ThorLabs SC10) separately for white light and fluorescence lamp synchronized with a cooled charge-coupled device (CCD) from Hamamatsu C4742-95 / C8484-03G02 or Scion corporation CFW1612M. The devices were controlled by custom made scripts using either Hamamatsu Wasabi or Micromanager interfaces. In order to obtain a large field of view, we used objectives with magnifications of 4x (0.13 NA Phase, Olympus) and 10x (0.25 NA Phase, Olympus). For phase contrast imaging, we also used an incubator microscope – SANYO MCOK-5M, integrated with a 10x phase contrast objective, and a Sentech XGA color CCD to scan multiple samples illuminated by LED. The unit is controlled by an MTR-4000 software. For laser ablation experiments, we used a Leica TCS SP5 confocal system equipped with Z-galvo stage and multiphoton infrared femtosecond pulsed laser (Coherent Chameleon Ultra II). The SP5 setup is mounted with an inverted Leica DMI6000 microscope, PMT and HyD detectors that are controlled by LAS AF acquisition interface. When the monolayer starts to exhibit *flows* after blebbistatin washout, cells within an area of ≈ 150 *μ*m × 150 *μ*m were removed by ablation. The ablation was performed with smaller Region of Interests (ROI) within the ≈ 150 *μ*m × 150 *μ*m area to avoid damaging the neighboring cells. The ablation was performed with Leica 63x (1.4 NA) oil objective at 800 nm wavelength and ≈ 4300 mW laser power. All experiments were performed at 37°C and images were acquired at 5 min, 10 min or 20 min interval between frames. In order to prevent evaporation of media due to long-term acquisitions (48 h – 60 h), samples were covered with a glass Petri dish or by adding 2 ml of mineral oil.

### Velocity Field and Correlation Functions

For all the analysis, the time point at which the monolayer reached complete confluency was assigned as time zero (*t*_0_). In order to obtain velocity field **v**(*x, y, t*) in the monolayer, Particle Image Velocimetry (PIV) was performed using the open source software PIVlab [68]. Specifically, the velocity components (*v*_*x*_, *v*_*y*_) at particular time *t* were obtained by using Direct Cross Correlation (DCC) algorithm on the monolayer images at times *t* and *t* + Δ*t* for interrogation window-size and step-size of 64 pixels and 32 pixels, respectively. The grid positions where no appropriate velocities could be obtained from the DCC algorithm were provided with velocity values via interpolation from the neigbouring grid points. Divergence 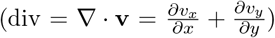 for the velocity field was calculated numerically using finite difference derivatives on the square grid plots. Likewise, the curl of the velocity field, curl 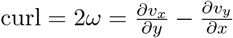, where *ω* is the vorticity, is also numerically calculated. Thus for each of *v*_*x*_, *v*_*y*_ and div, we obtain a *N*_*y*_ × *N*_*x*_ × *N*_*t*_ size matrix, where *N*_*x*_ and *N*_*y*_ are the number of grid points in the *x* and *y* directions, respectively, and *N*_*t*_ is the number of time frames. The spatio-temporal velocities and divergence thus obtained were further processed to calculate correlation functions in order to identify the underlying length and time scales in the following manner.

The velocity correlation function was obtained as:

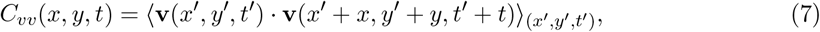

where the averaging was done over the space-time grid points *x*^′^, *y*^′^, and *t*^′^. The full correlation function was then azimuthally averaged to obtained *C*_*vv*_(*r, t*), where 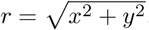. The spatial correlation function *C*_*vv*_(*r*) = *C*_*vv*_(*r*, 0), whereas the temporal correlation function *C*_*vv*_(*t*) = *C*_*vv*_(0, *t*). Please note that for convenience we use the same expression *C*_*vv*_ for the different variants of the correlation function. Correlation functions for divergence were similarly obtained starting from the full expression

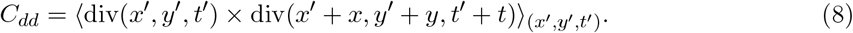

Length and time scales for pulsations and flows were extracted from the minima of the plots. We checked that the length and time values were consistent with actual motions measured on movies.

### Winding Number

From visual observations of the monolayer time-lapse movies, we noticed that pulsations in the tissue had focal points that corresponded to the locations where the velocities were lower than the surrounding regions. We formally quantified the location of these singular points by calculating the winding number for the velocity field on the smallest closed cell of the underlying rectangular grid as follows.

As shown in Figure S1a, there are four nodes *i* ∈ {1, 2, 3, 4} with velocity **v**_*i*_ at the respective node. The change in the angle between the velocity vector at a given node and the next was estimated as

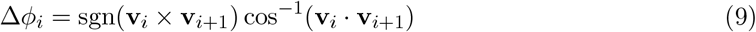

The expression sgn(**v**_*i*_ × **v**_*i*+1_) determines if the sense of rotation from *i* to *i* + 1 is clockwise or anti-clockwise. The second expression cos^−1^(**v**_*i*_ · **v**_*i*+1_) gives the smallest angle between *i* and *i* + 1. The total winding number *k* over the contour is

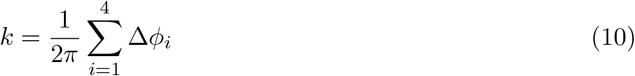

where the loop is traversed in an anti-clockwise sense (Figure S1b). If a singular point is present inside this loop then *k* takes an integer value – else it is zero. The singular points and their nature were further confirmed by plotting the streamlines of velocity fields. They broadly correspond to two main scenarios: (i) vorticity or source/sink (*k* = +1) and (ii) saddle point (*k* = −1). We found that the focal points of pulsation typically had a strength of *k* = +1 and corresponded either to a source ∇ · **v** *>* 0 or a sink ∇ · **v** *<* 0.

### Additional Details for the Computational Model

The active vertex model with cell motility is defined using Eqs. 1-6 of the main paper. In this model, both the absolute and the relative values of *ξ* (polarity velocity alignment) and *D*_*r*_ (polarity rotational diffusion) are crucial in dictating the nature of collective migration modes in the tissue. High values of *ξ* promote polarity alignment in a particular direction and hence long range flows. On the other hand, high values of *D*_*r*_ promote fast and short range oscillations. When the relative values of *D*_*r*_ and *ξ* are well-balanced, we find temporally periodic and spatially oscillatory migration patterns of the cells that mimic the experimental dynamics. In the friction grid experiments, we mimicked the grid-like patterns that were created in the experiments with regions of high and low substrate friction. This was achieved in the simulations by increasing the value of *η* in the fibronectin patch relative to the value outside of the patch. While modifying the value of *η*, we ensured that the magnitude of the motile force *ηv*_0_ approximately remained the same both inside and outside of the fibronectin patch. Tissue dynamics in typical simulations is reported along with experimental movies (Movies 12-15.)

The time-lapse images from the simulations have dimension of 1000 × 1000 px^2^ and are collected approximately after every 50 simulation time-steps. As in the experiments, the images from simulations are analyzed using PIV to get the velocity field [68, 69]. This velocity field is smoothened using Gaussian filter of size 2 px using pivmat to reduce the noise arising from short-range spatial noise in the movements of individual cells [69]. The divergence of this smoothed velocity field is obtained numerically using finite difference method [69].

#### Parameter values for the vertex model

A detailed list of the parameters used in the system and their corresponding non-dimensionalisation with respect to some length (*l*_0_), time (*t*_0_) and force (*f*_0_) scales is shown in the accompanying table. The force scale *f*_0_ is arbitrary since the force parameter appears on both left and right hand sides of the dynamical equation and hence need not be chosen. For the simulations we choose the non-dimensionalised simulation time-step Δ*t*^′^ = 1. The most obvious choice of the length scale *l*_0_ is related to the cell-dimension 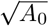. Based on these parameters, we can also extract additional length (*L*) and time (*τ*) scales.

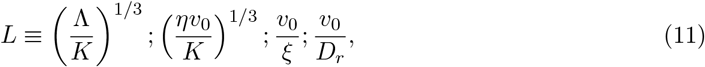

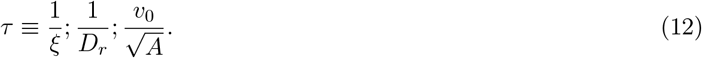

#### Comparison of simulation numbers with experimental values

There is an excellent qualitative correspondence between the experimental findings (Figures 1, 2, S5, 3a-b and S6b) and the simulation results (Figures 5 and S8). Below, we make quantitative connections between the experimental and simulation results by first linking the length and time-scales for plots of Figure 5 with the experimental units.

The simulation velocity field of cells was quantified using PIV on images of 1000×1000 px^2^, sampled after Δ*t*^′^_*f*_ ≈ 50Δ*t*^′^, where Δ*t*^′^ is the simulation time-step and ^′^ denotes non-dimensionalised value. The distance *R* in the correlation function of Figure 5d is reported in px units and time *t* in terms of the duration between two frames (Δ*t*^′^_*f*_). Since there are *N* = 3060 cells in the 10^6^ px^2^ image, each cell length *l*_cell_ corresponds to

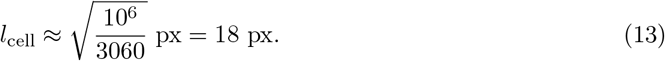

The minima of the correlation function for Figure 5d is between 150 − 200 px. Hence, the collective pulsatile movement occurs approximately on the scale of 8 − 11 cell lengths. This number is similar to the experimentally observed collective length-scale of pulsation. Moreover, by noting the *l*_cell_ ≈ 20 *μ*m, we also estimate that

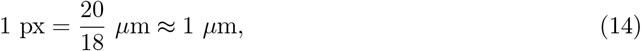

thus giving a correlation length of approximately 150 − 200 *μ*m, which is consistent with the experimental findings. This also provides the experimental measure for distance *R* in Figure 5. Going along these lines, we now make an estimate of the length scale (*l*_0_) that is implicit in the simulations. From Table 1, the preferred area of a single cell 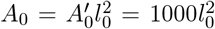. Taking *A*_0_ ≈ 20^2^ *μ*m^2^, we obtain

**Table 1.**
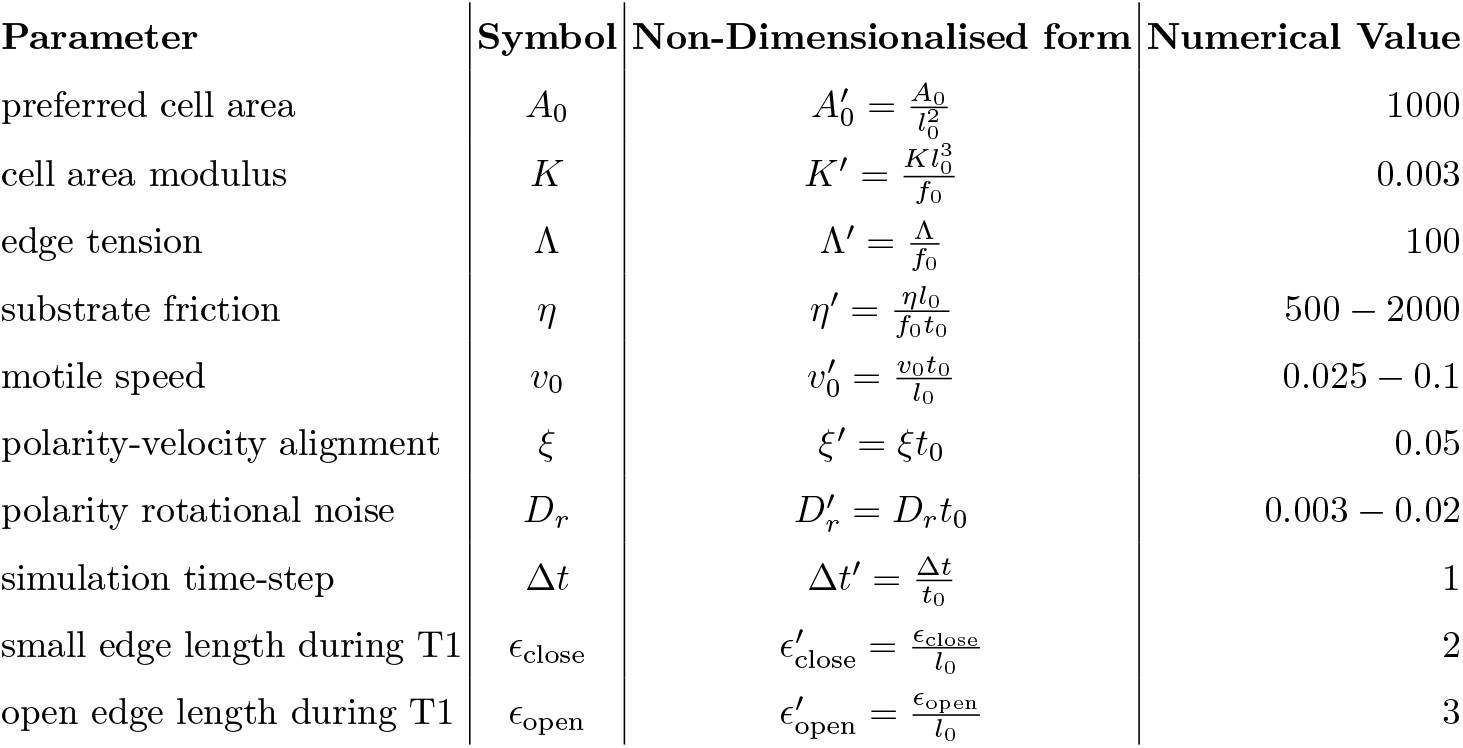
Parameters for the vertex model

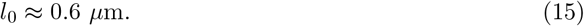

The minima for the correlation function in Figure S8c for the control case is at around 20Δ*t*^′^_*f*_ (frame sampling). This corresponds to a period of ≈ 40Δ*t*^′^_*f*_ ≈ 40 × 50Δ*t*^′^ = 2000Δ*t*^′^, as the duration between two frames is around 50Δ*t*^′^. Since the experimentally observed pulsation time-period is ≈ 5 h, we estimate

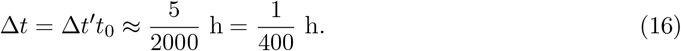

Since, we used Δ*t*^′^ = 1, the value of Δ*t* is also the same as the time-scale *t*_0_ that is implicit in the simulations (Table 1). Also, the time difference between two frames of simulation time-lapse images is

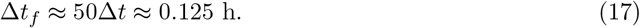

This is the experimental measure for the time in Figure 5. We check the consistency of this number as follows. In simulation units, the value of cell motility 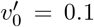. From Table 1 we can note that

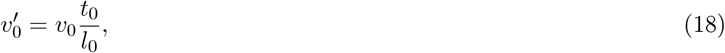

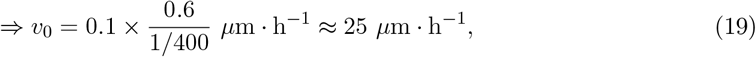

where we have used the estimates of *l*_0_ and *t*_0_ made above. This number is consistent with the experimental findings for cell speed.

### Velocity-polarisation correlation for migrating cells

The quantification for the alignment of lamellipodia direction with the local flows is shown in Figures 4e, S7f and S7g-h. This quantification is obtained as follows. We obtain the velocity field of the flow domain and cell contour separately and plot them together. The velocity field is obtained from the PIV of the domain using phase contrast images. From this, we compute the mean flow direction for the domain. For the lamellipodia direction, since cell direction is set by the leading front, we obtain the effective cell movement direction as a representative of the lamellipodia direction. Individual cells are tracked after blebbistatin washout over a duration of approximately 12 hours, and the cells are outlined.

We then obtain the PIV of the cell contour which gives the vectors along the cell contour during cell movement. From this, we get the mean direction/angle of cell movement. This angle is used to represent the lamellipodia direction. We visually verified that the lamellipodia are always at the cell front in the direction of cell movement.

We also used another method to obtain the polarisation of a cell as defined in Figure S2. As described in the previous paragraph, we already have tracked cells with outlines. We then obtain the centroid of the cell for every time-step *t* as follows:

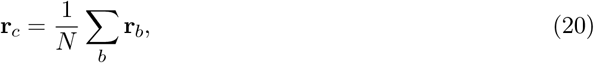

where *b* is the index, respectively, corresponding to the *N* boundary pixels. The polarity of the cell is then defined in terms of its nematic tensor :

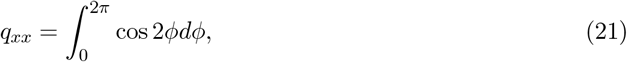

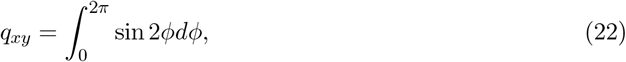

where the angle *ϕ* is defined between *x* axis and the line connecting the centroid and the boundary pixel as shown in Figure S2a. The polarity can then be defined as 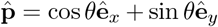, where *θ* = tan^−1^(*q*_*xy*_*/q*_*xx*_). The direction of cell velocity 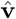 is obtained from the instantaneous displacement of the centroid between subsequent time steps. The correlation between the polarity and velocity direction thus defined is calculated by getting the frequency distribution over all time steps of the quantity

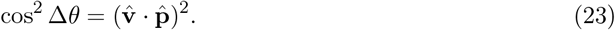

The combined histogram for this quantity from multiple cases is obtained in Figure S2. The distribution is skewed towards cos^2^ Δ*θ* = 1, i.e., Δ*θ*, the angle between the velocity and polarity is systematically biased towards smaller angles (Figure S2b).

We also obtained the connection between cell polarity, defined in terms of collective shape anisotropy, and cell velocity across different experimental conditions. We first acquired polarity field 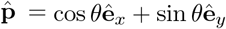 of a group of cells through cell shape anisotropy as calculated from the structure tensor of the time-lapse images for fibronectin grid and blebbistatin washout conditions. The actual analysis was performed using the OrientionJ plugin of FIJI [70]. The velocity field **v** of the cells was already obtained for these time-lapse images using PIVlab as discussed in the main paper. The polarity and velocity were both sampled on a 32 px × 32 px grid with indices *i, j* across the tissue domain examined for each of the experimental condition. To make an esimate of the degree of alignment between polarity 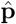 and velocity orientation 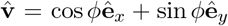, we calculated the cross-correlation

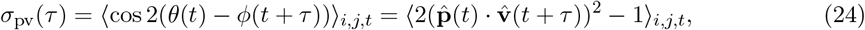

where the average was done over the the spatial grid points *i, j* for all time-frames *t*. Note that, as in the earlier paragraph, *σ*_pv_ was defined to satisfy 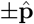 equivalence for cell shape anisotropy. We found that the value of *σ*_pv_(0) was approximately −0.02 and 0.55, respectively, for grid and blebbistatin experiments.

## QUANTIFICATION AND STATISTICAL ANALYSIS

Graph plotting and statistical analysis were performed using GraphPad Prism 7 software. Kruskal-Wallis test was performed to test the significance between control and different conditions in Figure S4d-i. The statistical significance is indicated as * p < 0.05. The number of experimental repeats is mentioned in the figure caption and the data is expressed as Mean value ± Standard deviation.

