## Supplementary Figures for "Pulsations and flows in tissues: two collective dynamics with simple cellular rules"

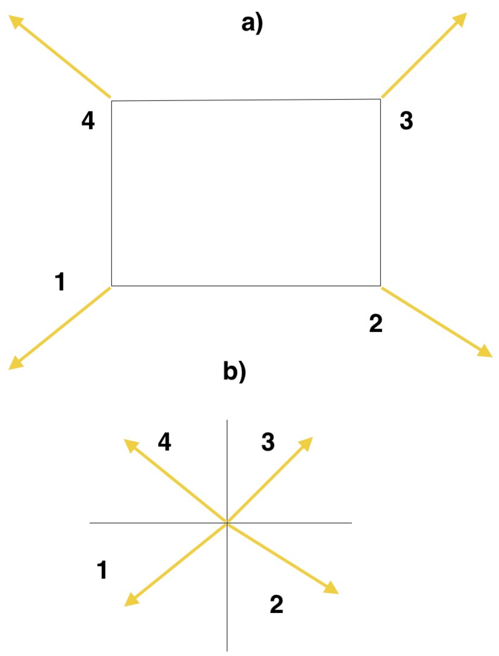


**Figure S1**: **Winding number calculation, related to Figures 1, 2, 3 and 5.** (a) Calculation of winding number for the velocity field over the smallest cell of the PIV grid as described in Winding number section in STAR Methods. (b) A clearer illustration of the angles between velocity vectors on subsequent grid points arranged in an anticlockwise sense. See the topological defect panels in the associated Figures 1, 2, 3, 5, S5, S6, S7 and S8.

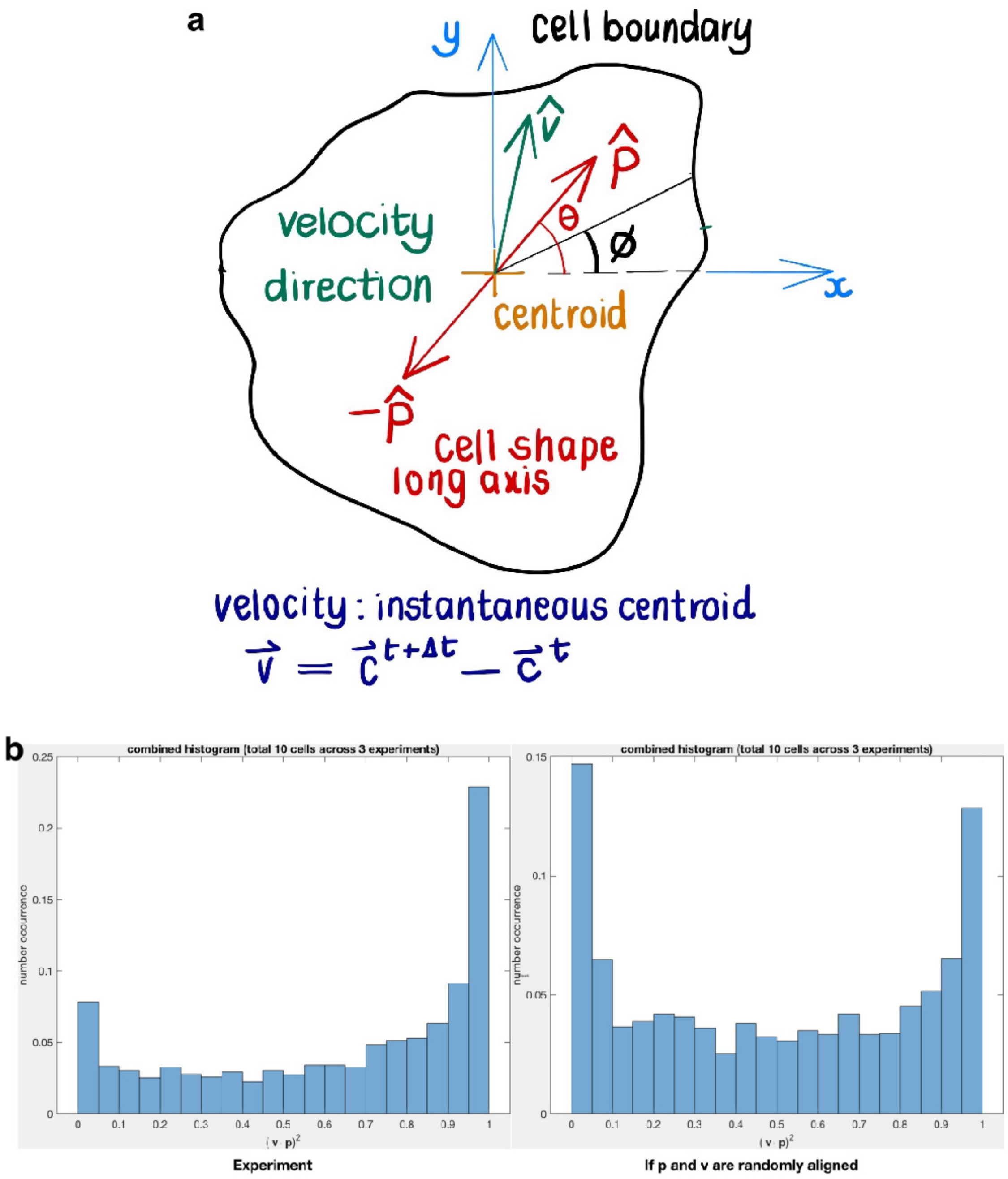


**Figure S2**: **Nematic order parameter from cell contour, related to Figure 4.** (a) The notation used to calculate polarisation and velocity of individually tracked cells. (b) Histogram is plotted for the quantity $\cos^2 \Delta\theta = (\hat{\mathbf{v}} \cdot \hat{\mathbf{p}})^2$ over a duration of $10$ h at $15$ min intervals for $10$ cells from different blebbistatin washout experiments (N = 3). A similar histogram is also plotted when $\Delta\theta$ for the same number of experimental data points, but now randomly sampled uniformly from $[0, \pi/2]$. This histogram is more symmetric for values at $\cos^2 \Delta\theta = 0$ and $\cos^2 \Delta\theta = 1$, when compared to its experimental counterpart. See associated Figure 4 and Figure S7. For details see Velocity-polarisation correlation for migrating cells section in STAR Methods.

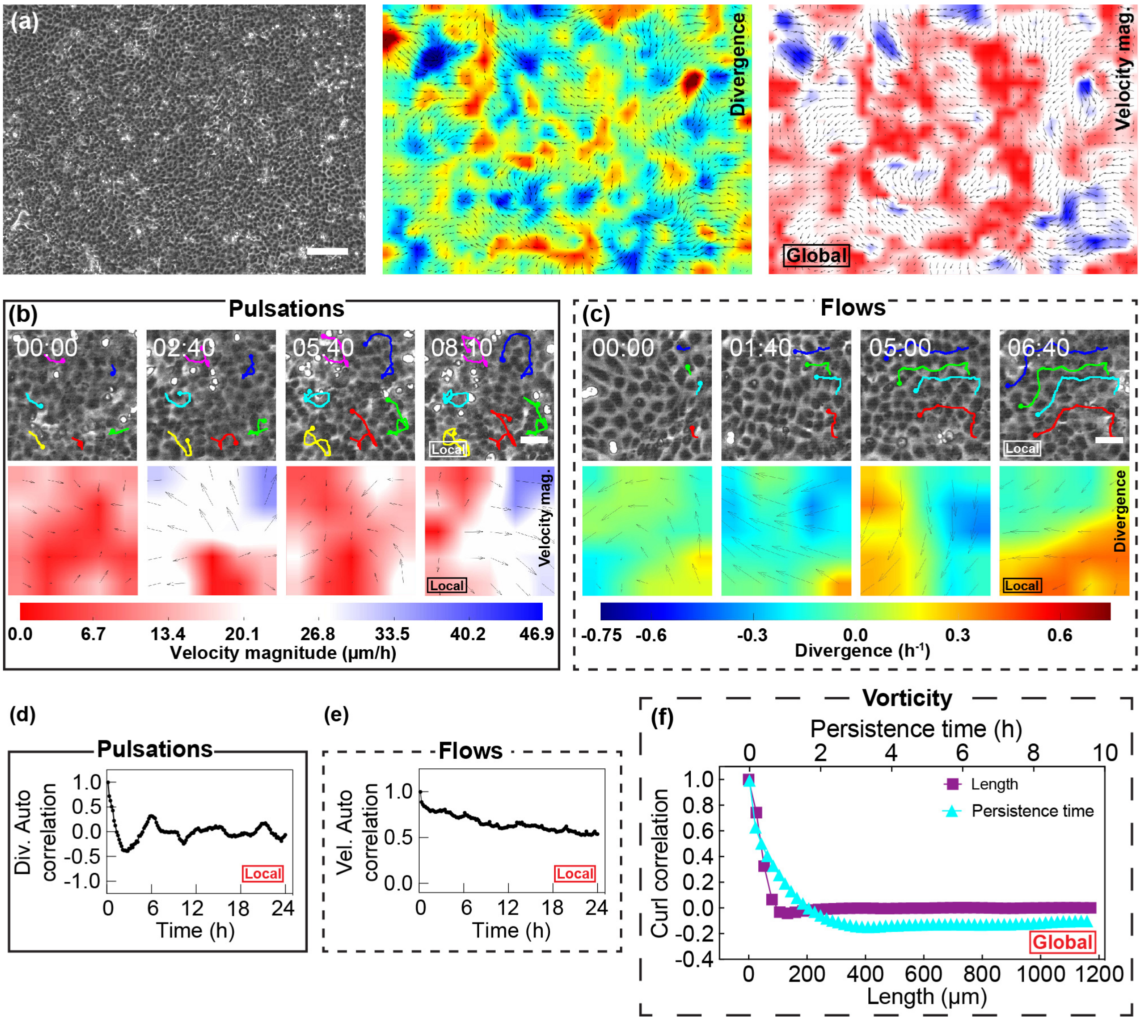


**Figure S3**: **Characteristics of pulsations and flows in monolayer, related to Figure 1.** (a) Snapshot of a confluent MDCK monolayer in phase contrast and the corresponding divergence and velocity magnitude colour maps. Scale bar, $200\ \mu m$. These snapshots correspond to the same experiment and data shown in Figure 1. In (b) and (c): first row shows the raw data with tracking for pulsation (b) and flows (c); second row shows the velocity magnitude for pulsations (b) and divergence for flows (c). (b) and (c) correspond to the data shown in Figure 1a and 1d respectively. Scale bar $50\ \mu m$. Time is in hh:mm. In those panels indicating velocity magnitude, red and blue indicate low and high velocities respectively; similarly, in those panels indicating the nature of the divergence field, blue indicates contraction, and red indicates expansion. The numerical values are specified in the corresponding color bars. (d) and (e) show the autocorrelation of divergence and velocity magnitude respectively, corresponding to Figure 1b and 1c. (f) represents the global characteristics of the monolayer and shows the vorticity correlation for distance and time over $48$ h, for the same experiment shown in Figure 1c and 1f.

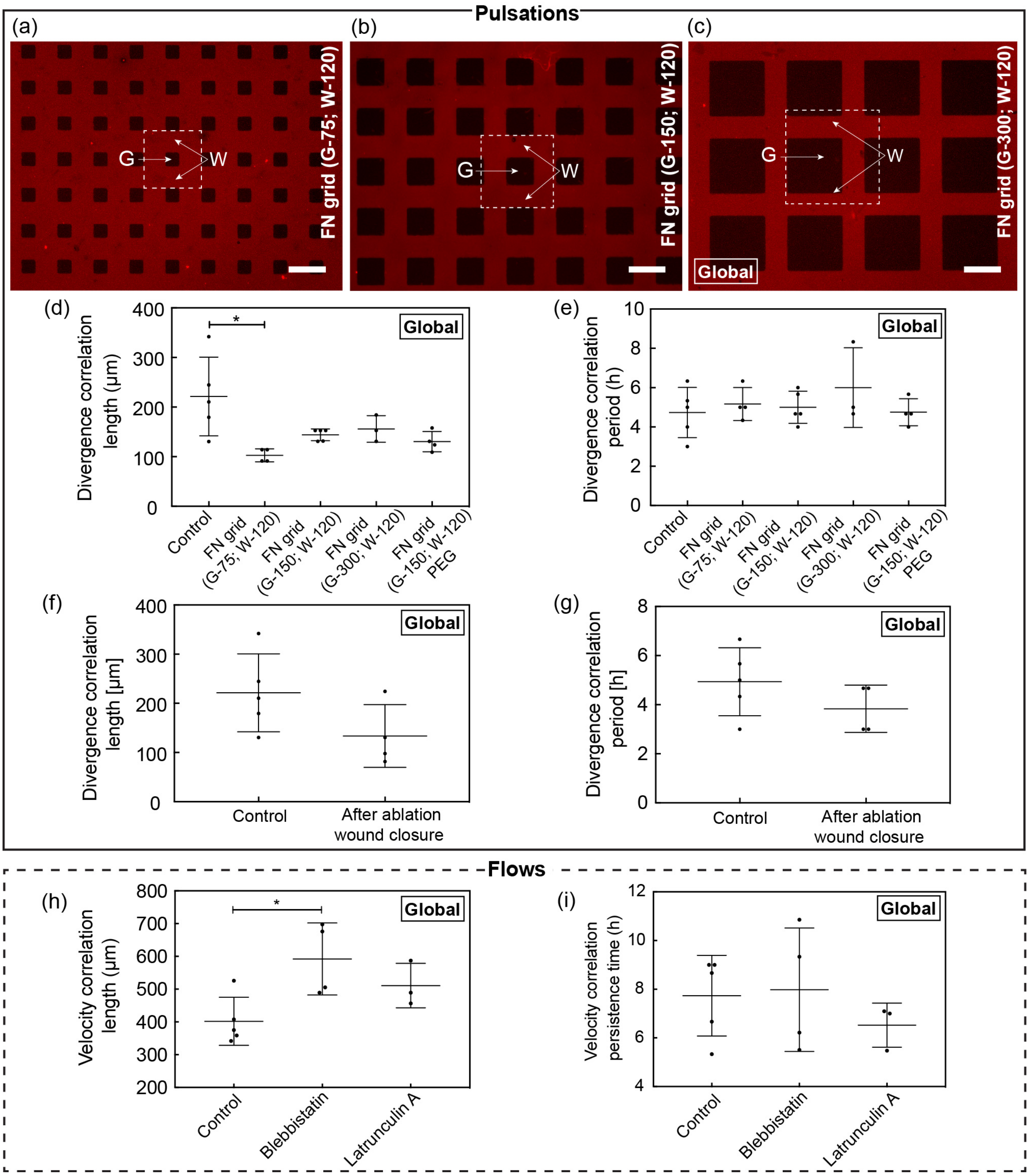
Pulsations
(a)
(b)
(c)
G
W
FN grid (G-75; W-120)
FN grid (G-150; W-120)
FN grid (G-300; W-120)
Global
(d)
Divergence correlation length (μm)
*
Control
FN grid (G-75; W-120)
FN grid (G-150; W-120)
FN grid (G-300; W-120)
FN grid (G-150; W-120) PEG
(e)
Divergence correlation period (h)
(f)
Divergence correlation length [μm]
After ablation wound closure
(g)
Divergence correlation period [h]
Flows
(h)
Velocity correlation length (μm)
Blebbistatin
Latrunculin A
(i)
Velocity correlation persistence time (h)

**Figure S4**: **Different fibronectin (FN) grid and flow conditions, related to Figures 2 and 3.** (a), (b) and (c) show the FN grids with different sizes. Each grid unit, highlighted in dotted lines in (a), (b) and (c), has a non-FN area or a gap (G) shown in black, and a surrounding FN region shown in red. In all conditions, the FN width (W) is maintained constant at $120\ \mu\mathrm{m}$ while the gap (G) is varied: $75\ \mu\mathrm{m}$ in (a); $150\ \mu\mathrm{m}$ in (b); and $300\ \mu\mathrm{m}$ in (c). Scale bar, $200\ \mu\mathrm{m}$. See the associated Figures 2 and S5. (d), (e), (f), (g), (h) and (i) represent the global characteristics of the monolayer and all the data are expressed as Mean value $\pm$ Standard deviation. (d) and (e) show the mean divergence correlation lengths and periods, respectively for different FN grid conditions, obtained over 48 h. In (d), * indicates a significant difference ($p < 0.05$). While the smallest FN grid (G-75; W-120) shows a significant difference from the control, other FN grid conditions show smaller variations between the experiments as opposed to control which shows a larger variation range. This emphasizes the effect of different FN grid conditions in regulating the correlation length. The divergence correlation period shown in (e) remains unaffected. In both (d) and (e), each dot corresponds to an individual experiment (N) where the N for each condition is as follows: Control (N = 5); FN grid (G-75; W-120) (N = 4); FN grid (G-150; W-120) (N = 5); FN grid (G-300; W-120) (N = 3); and FN grid (G-150; W-120) with PEG (N = 4). See the associated Figures 2 and S5. (f) and (g) show the divergence correlation lengths and periods respectively, of the monolayer following ablation wound closure after the blebbistatin washout, obtained over $24$ h. The difference between the conditions in (f) is statistically non-significant and shows a diverse spread in the correlation length values similar to control. The observed shift towards lower correlation length for the ablation wound closure could be due to the increased cell density. The divergence correlation period shown in (g) remains almost unaffected. In both (f) and (g), each dot corresponds to an individual experiment (N) where N for each condition is as follows: control (N = 5); After ablation wound closure (N = 4). See the associated Figures 3 and S6. (h) and (i) show the mean velocity correlation lengths and times, respectively for the different flow conditions, obtained over 36 h. In (h), * indicates a significant difference ($p < 0.05$) for the blebbistatin condition and the latrunculin A condition shows a shift towards higher correlation lengths, compared to control. The persistence time shown in (i) remains almost unaffected. In both (h) and (i), each dot corresponds to an individual experiment (N) where the N for each condition is as follows: Control (N = 5); Blebbistatin (N = 4); and Latrunculin A (N = 3). See the associated Figures 3 and S6.

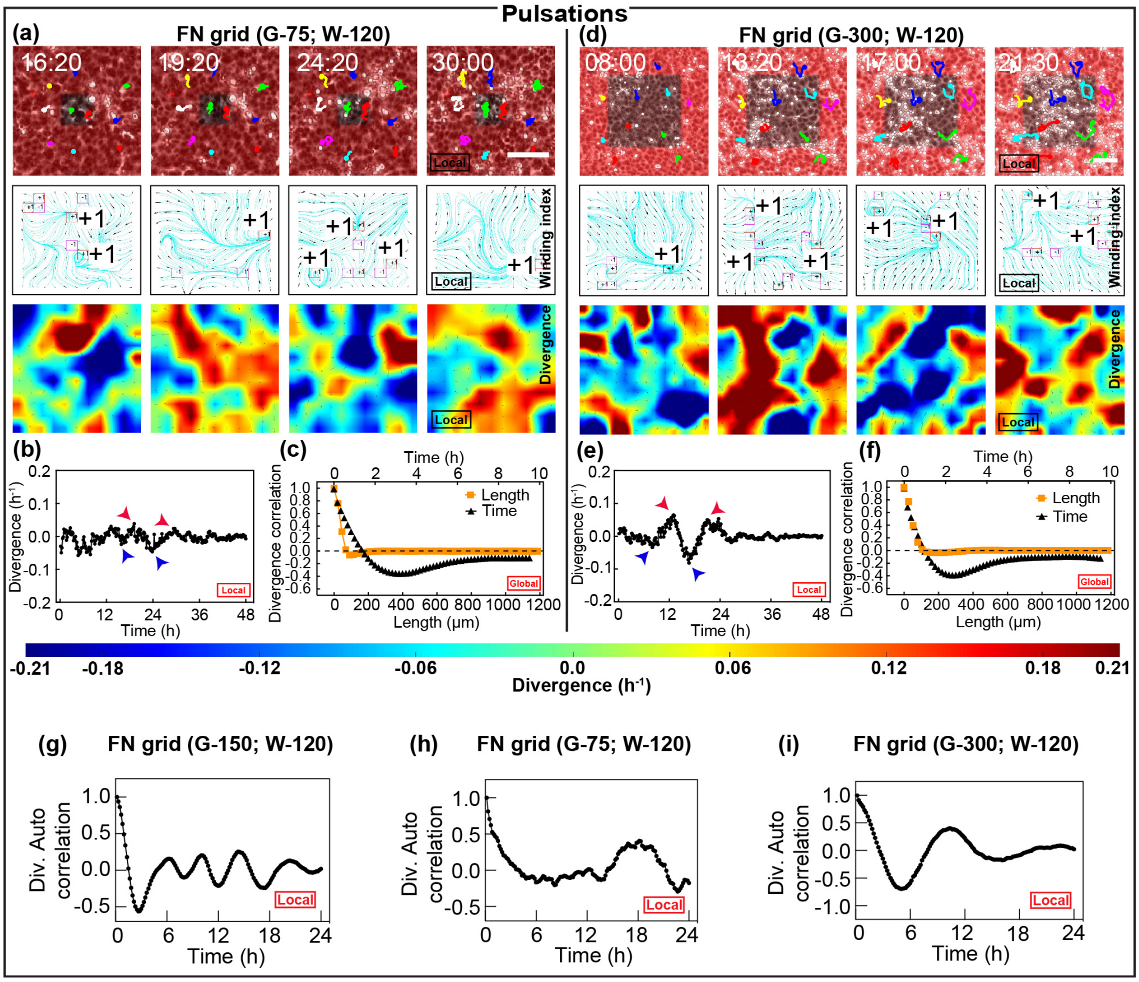


**Figure S5**: **Variation in Fibronectin (FN) grid sizes alter pulsation characteristics, related to Figure 2.** (a)-(c) and (d)-(f) correspond to smaller ($G = 75\ \mu\mathrm{m}$) and larger ($G = 300\ \mu\mathrm{m}$) FN grid conditions respectively. The first row of (a) and (d) show phase contrast images of cells on a single FN grid unit where cell trajectories are highlighted by tracking. The second and third rows show the topological defects and the divergence, respectively. The colour bar shows the scale for divergence where blue indicate contraction and red indicates expansion. Scale bar, $100\ \mu\mathrm{m}$ and time is in hh:mm. See also the associated Movies 3 and 5. (b) and (e) show the mean divergence plots over 48 h, and arrows indicate the snapshots shown in the domains (a) and (d) respectively. (c) and (f) show the divergence correlation plots in space and time for the entire region. (g), (h) and (i) show the divergence autocorrelation for different FN grid conditions, corresponding to Figures 2b, S5b and S5e, respectively. See also the associated Figure 2.

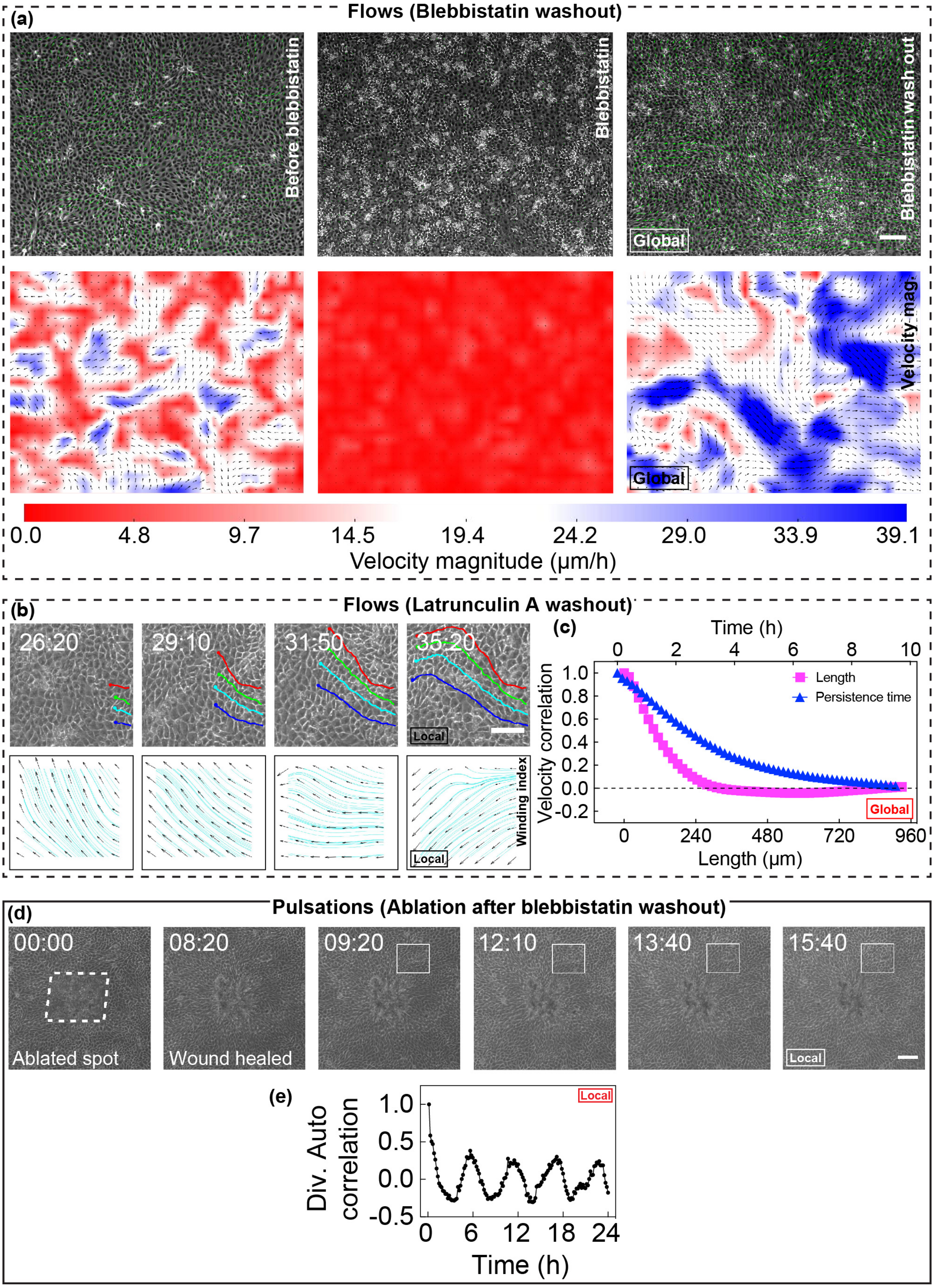
(a)
Flows (Blebbistatin washout)
Before blebbistatin
Blebbistatin
Blebbistatin wash out
Global
Velocity mag.
Global
0.0
4.8
9.7
14.5
19.4
24.2
29.0
33.9
39.1
Velocity magnitude (μm/h)
(b)
Flows (Latrunculin A washout)
26:20
29:10
31:50
35:20
Local
Winding index
Local
(c)
Time (h)
0
2
4
6
8
10
Velocity correlation
1.0
0.8
0.6
0.4
0.2
0.0
-0.2
Length
Persistence time
Global
0
240
480
720
960
Length (μm)
(d)
Pulsations (Ablation after blebbistatin washout)
00:00
Ablated spot
08:20
Wound healed
09:20
12:10
13:40
15:40
Local
(e)
Local
Div. Auto correlation
1.0
0.5
0.0
-0.5
0
6
12
18
24
Time (h)

**Figure S6**: **Emergence of long range flows after Blebbistatin and Latrunculin A washouts, related to Figure 3.** (a) shows snapshots of monolayer during the resetting of myosin activity. The three snapshots represent the states before incubation with blebbistatin, in the presence of blebbistatin, and after washing out blebbistatin, respectively. The first row shows the phase contrast images of cells superimposed with velocity vectors in green. The second row shows velocity magnitude where red and blue indicate low and high velocities, respectively. The velocity is very low in the presence of blebbistatin, and flow patterns emerge after blebbistatin washout. Scale bar $200\ \mu\mathrm{m}$. These snapshots correspond to the experiment and data shown in Figures 3a and 3b. (b)-(c) Transition to flows using Latrunculin A. The first row of (b) shows phase contrast images of a domain of cells where tracking highlights the cell trajectories. Correspondingly, the second row shows the absence of topological defects. Scale bar $100\ \mu\mathrm{m}$ and time in hh:mm. (c) shows persistence in space and time for velocity correlation obtained over the entire region surrounding the domain shown in (b). (d) Ablation performed on flows resulting from resetting myosin activity. (d) shows a snapshot of phase contrast images of the ablated spot and the surrounding area. Wound due to ablation is highlighted by a dotted line in the first time point which eventually heals (second time point). In the consecutive time points, a smaller box is drawn to highlight a region exhibiting pulsatile behaviour. Scale bar $100\ \mu\mathrm{m}$. Time is in hh:mm. (e) shows the divergence autocorrelation following the ablation wound closure for the pulsatile domain (small box) in (d), where the time periodicity is apparent. (d) and (e) correspond to the same experiment and data shown in Figures 3c, 3d and 3e. See the associated Movie 9.

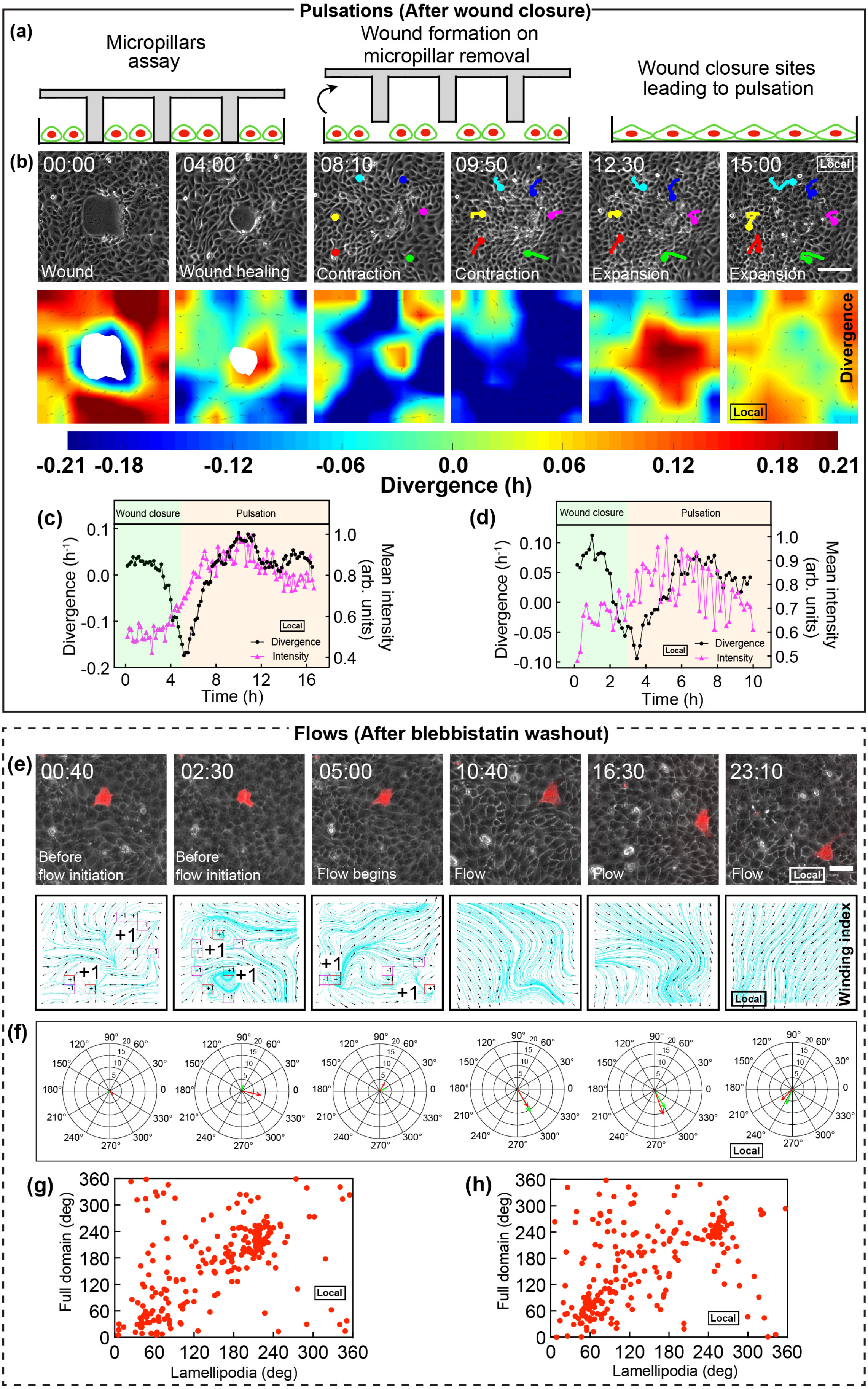

Pulsations (After wound closure)
(a)
Micropillars assay
Wound formation on micropillar removal
Wound closure sites leading to pulsation
(b)
00:00
04:00
08:10
09:50
12:30
15:00
Local
Wound
Wound healing
Contraction
Contraction
Expansion
Expansion
Divergence
Local
-0.21 -0.18 -0.12 -0.06 0.0 0.06 0.12 0.18 0.21
Divergence (h)
(c)
Wound closure
Pulsation
Divergence (h-1)
Mean intensity (arb. units)
Local
Divergence
Intensity
Time (h)
(d)
Wound closure
Pulsation
Divergence (h-1)
Mean intensity (arb. units)
Local
Divergence
Intensity
Time (h)
Flows (After blebbistatin washout)
(e)
00:40
02:30
05:00
10:40
16:30
23:10
Before flow initiation
Before flow initiation
Flow begins
Flow
Flow
Flow
Local
+1
Winding index
Local
(f)
Local
(g)
Full domain (deg)
Lamellipodia (deg)
Local
(h)
Full domain (deg)
Lamellipodia (deg)
Local

**Figure S7**: **Micropillar and mosaic experiments for investigating the molecular actors behind pulsations and flows, respectively, related to Figure 4.** (a)-(b) Micropillar assay showing pulsatile behavior after wound closure. (a) Schematic of the micropillar assay. The monolayer was grown in the presence of PDMS micropillars which on removal leads to wounds for the monolayer to close. (b) Wound closure leading to pulsations. First and second rows show snapshots of phase contrast images and the associated divergence maps, respectively, corresponding to Figure 4a and 4b. In both rows, the first and second time points represent the wound and its eventual closure, respectively. The last four time points represent the pulsation that follows wound closure. Tracking in the first row highlights the back and forth movement of cells. The color bar indicates the divergence where blue represents contraction and red represents expansion. The snapshots with blue and red regions in second row correspond to the densification and diffusion of myosin in Figure 4a and 4b, at time points 12:40 and 15:10, respectively. Scale bar $100\ \mu\mathrm{m}$ and time is in hh:mm. (c) and (d) show the divergence and myosin intensity plots for two different pulsatile domains from two separate experiments. The green region represents the wound closure phase, and the pink region represents the pulsation phase. (e) and (f) show the orientation of lamellipodia aligning with the direction of flow. First two time points in (e) and (f) represent the phase before flow initiation; the transition is shown by the third time point; and the last three time points represent the flow phase. (e) The first row shows the phase contrast images of MDCK-E-cadherin-GFP cells where a cell transiently transfected with LifeAct-mCherry is shown in red to demonstrate the orientation of lamellipodia before and after initiation of flow. The second row shows the topological defects. The presence and absence of topological defects before (first two snapshots) and after (last three snapshots) initiation of flow demonstrates the transition to flow behaviour. Scale bar $50\ \mu\mathrm{m}$ and time is in hh:mm. (f) Plot showing the alignment of lamellipodia (obtained from the effective cell direction - see Velocity-polarisation correlation for migrating cell section in the STAR Methods) and mean direction of the flow field (obtained for the snapshots in (e)). The direction is given by the distribution of angles corresponding to the 2D space of images shown in (e). Velocity magnitude ($\mu\mathrm{m}$.h-1) is given by the concentric circles. The green arrow corresponds to the mean direction of flow field (indicated by streamlines in (e)), and the red arrow corresponds to the mean orientation angle of lamellipodia (shown by the red cell in (e)). (e) and (f) correspond to the same experiment and data shown in Figures 4d and 4e. (g) and (h) Plots showing the linear tendency between lamellipodia orientation and the mean flow direction for two different domains from two separate experiments. (see Velocity-polarisation correlation for migrating cells section in the STAR methods).

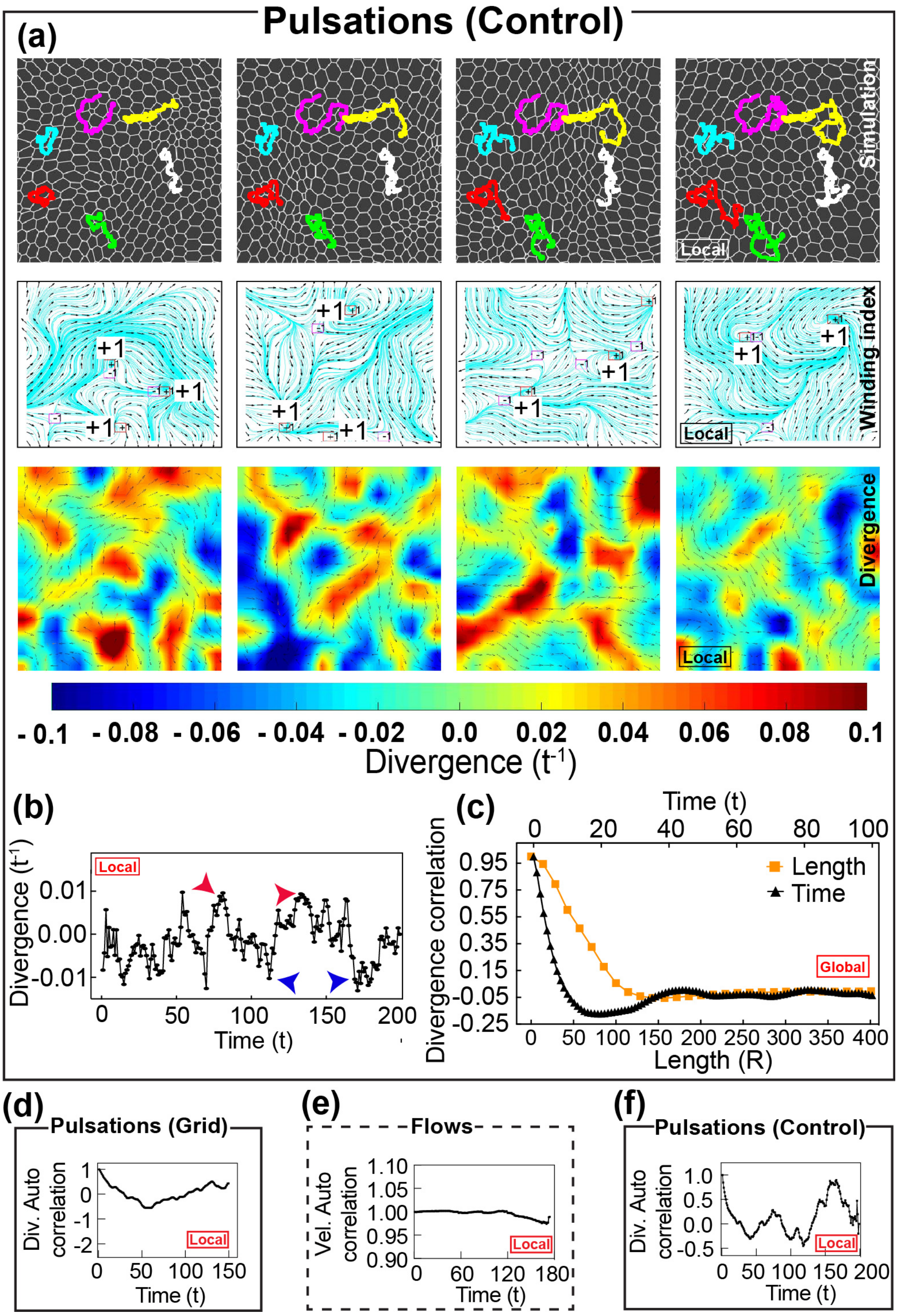


**Figure S8**: **Simulation of pulsations using active vertex model, related to Figure 5.** (a)-(c) correspond to their experimental counterparts in Figure 1 (a)-(c). (a) shows a domain of the simulated monolayer, and the tracking highlights the pulsatile movements of cells in the first row. Topological defects and divergence are shown in the second and third rows, respectively. Contraction and expansion in the color-coded divergence maps are represented by blue and red colors, respectively. See the associated Movie 12. (b) shows divergence plotted as a function of time for the domain shown in (a). The periodicity of the divergence is shown by the autocorrelation in (f). (c) Divergence correlation function in space and time for the entire region surrounding the domain shown in (a) shows the emergence of length and time scale similar to experiments. See Comparison of simulation numbers with experimental values section in STAR Methods for discussion on experimental units of time and length. (d), (e) and (f) show the autocorrelation for simulated grid condition, flows and control condition, corresponding to Figures 5c, 5f and S8b.

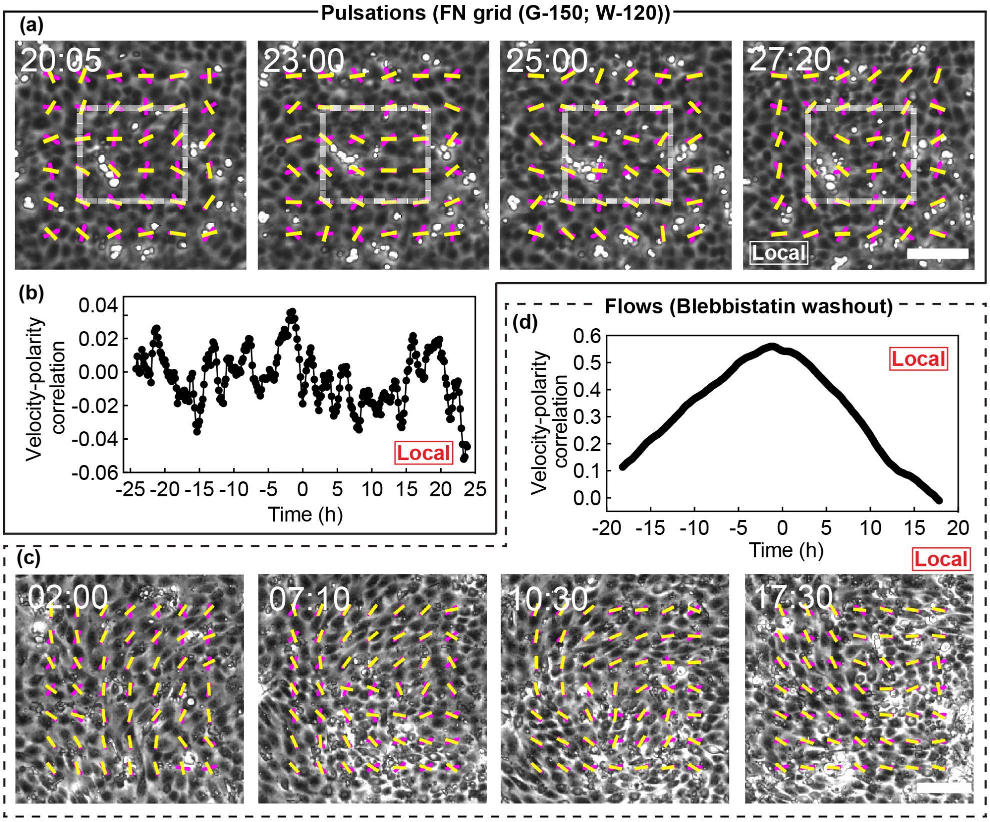


**Figure S9**: **Velocity and polarity alignment in pulsations and flows, related to Figures 2 and 3.** (a)-(b) show pulsations data and (c)-(d) show flow data. The images in (a) correspond to the same images of FN grid condition ($G = 150\ \mu\mathrm{m}$, outlined in grey) shown in Figure 2a and the images in (c) correspond to the same images of blebbistatin washout condition shown in Figure 3a. The magenta and yellow lines in (a) and (c) represent the velocity direction and polarity, respectively, and illustrate the alignment between them. The alignment is quantified as velocity-polarity correlation that is plotted in (b) and (d) for the domains shown in (a) and (c), respectively. The velocity-polarity correlation values are much higher in the flow condition (d), compared to the pulsations in grid condition (b). Scale bar is $100\ \mu\mathrm{m}$ and time is in hh:mm in (a) and (c).
