## Supplementary material for "Pulsations and flows in tissues: two collective dynamics with simple cellular rules": Resource table

**Resources table**

| REAGENT or RESOURCE | SOURCE | IDENTIFIER |
|---|---|---|
| Chemicals, peptides, and recombinant proteins | | |
| Dulbecco's Modified Eagle's Medium (DMEM) | Invitrogen | 31885-049 |
| Fetal Bovine Serum (FBS) | HyClone | 10309433 |
| Penicillin-Streptomycin | Invitrogen | 11548876 |
| Leibovitz L-15 medium | Invitrogen | 11540556 |
| Blebbistatin | Sigma Aldrich | B0560 |
| Latrunculin A | Sigma Aldrich | L5163 |
| PolyDimethyl Siloxane (PDMS) | Dow Corning | Sylgard 184 |
| SU-8 | MicroChem | - |
| Hydrogen Peroxide | Sigma Aldrich | 516813 |
| Sulphuric Acid | Sigma Aldrich | 258105 |
| (3-mercaptopropyl)trimethoxysilane | Fluorochem | S10475 |
| TRITC labelled Fibronectin | Cytoskeleton | FNR01 |
| HiLyte 488 labelled Fibronectin | Cytoskeleton | FNR02 |
| Phosphate Buffered Saline (PBS) | Invitrogen | 11530486 |
| PLL-g-PEG (poly-L-Lysine-grafted-PolyEthylene Glycol) | SuSoS | - |
| Pluronic acid | Sigma Aldrich | P2443 |
| Mineral oil | Sigma Aldrich | M8410 |
| Experimental models: Cell lines | | |
| MDCK-GFP-E-Cadherin | James W. Nelson lab., Stanford University | - |
| MDCK-mCherry-Actiin-GFP-Myosin | Roland Wedlich-Söldner lab., University of Münster | - |
| MDCK-GFP-Myosin | Shigenobu Yonemura lab., RIKEN | - |
| Software and algorithms | | |
| Fiji | Schindelin et al., 2012 | https://imagej.net/software/fiji/ |
| OrientationJ plugin, Fiji | Püspöki et al., 2016 | http://bigwww.epfl.ch/demo/orientation/ |
| MATLAB | - | https://se.mathworks.com/products/matlab.html |
| PIVlab, MATLAB | Thielicke et al., 2014 | https://se.mathworks.com/matlabcentral/fileexchange/27659-pivlab-particle-image-velocimetry-piv-tool-with-gui |
| Pivmat | Frédérick et al., 2017 | http://www.fast.u-psud.fr/pivmat/ |
